# In silico optimization of DNA codons in genes encoded by various strains of Ebola Virus

**DOI:** 10.1101/2023.09.24.559163

**Authors:** Anshu Mathuria, Mehak Ahmed, Indra Mani

## Abstract

Ebola hemorrhagic fever (HF) is a severe and often lethal disease that occurs in primates, including humans. It was first identified in 1976 during two outbreaks of fatal haemorrhagic fever in Central Africa. Due to its high fatality rate and lack of a widespread cure, Ebola HF poses a considerable challenge in treatment, making Ebola virus disease (EVD) one of the deadliest zoonotic diseases. The viral genome is ∼19kb long, linear, non-segmented, and negative single-stranded (-SS) RNA. The genome of the Ebola virus (EBOV) comprises seven genes, namely NP, GP, L, and VP (VP30, VP24, VP40, VP35), which encode nucleoprotein, glycoprotein, RNA polymerase, and viral proteins. By optimizing the DNA sequence through codon adaptation, we observed significant enhancements in the codon adaptation index (CAI) and the GC content compared to the wild-type strain. These findings demonstrate that optimized genes hold the potential for improved expression in the host organism without the production of truncated proteins. Further, these optimized genes can facilitate proper protein folding and function. In conclusion, these results have implications for vaccine production, as higher codon optimization enhances the expression of the genes, making them appropriate amounts for vaccine development.

## 1. Introduction

The Ebola virus (EBOV) genome is ∼19kb long, non-segmented, linear, and negative single-stranded (-SS) RNA (Jain et al., 2021). EBOV is a zoonotic filovirus that possesses an envelope. It has been classified into five species: *Zaire ebolavirus*, *Tai Forest ebolavirus* (formerly known as Cote d’Ivore), *Bundibugyo ebolavirus*, *Sudan ebolavirus*, and *Reston ebolavirus* (It is found in the Western Pacific region and is highly pathogenic in non-human primates) (Goeijenbier et al., 2014). The fruit bat is considered the animal reservoir of EBOV. Ebola hemorrhagic fever (Ebola HF) is transmitted to humans mainly through direct contact with animal tissues or body fluids. This includes activities such as processing and eating animal meat and consuming contaminated water and bat droppings. Infections are often associated with transmission in humans. EBOV is typically transferred through direct exposure of mucous membranes or breaks in the skin to infected body fluids (Kanapathipillai et al., 2014). Furthermore, two other genera are included in the Filoviridae family, *Cuevavirus* and *Marburgvirus* (http://www.ictvonline.org/virusTaxonomy.asp). The primary source of infection during an epidemic is coming in contact with either sick individuals or human corpses. However, fruit bats, which are asymptomatic carriers of the virus, are believed to be the natural reservoir. Isolates of filoviruses obtained from fruit bats exhibit a higher level of genetic diversity (Towner et al., 2009; Carroll et al., 2013). As reported by WHO on 21^st^ November 2014, the ongoing outbreak of Ebola HF resulted in a total of 15,351 confirmed or suspected cases, with 5,459 reported deaths. Among the countries most affected by the epidemic are Liberia, Guinea and Sierra Leone(http://apps.who.int/iris/bitstream/10665/144117/1/roadmapsitrep_21Nov2014_eng.pdf?ua=1).

The Ebola virus’s virion maintains a consistent diameter of approximately 80 nm, while its length varies within the range of 970 to 1200 nm. However, when grown in the cell culture, it exhibits a marked pleomorphism that can extend up to 14,000 nm (Geisbert and Jahrling 1995). The virion core contains an unsegmented, linear, negative-sense RNA molecule. This RNA is coiled and interacts with the nucleoprotein (NP), VP30, VP35, and polymerase (L) proteins. The nucleocapsid is helical and surrounded by a layer of specific glycoprotein spikes about 10 nm long. These glycoproteins have a crucial role in the virus’s infectivity by facilitating virus entry and contributing to its immunogenicity. They are targeted by immune cells and are therefore considered important in vaccine development. The viral matrix proteins VP40 and VP24 are situated between the outer envelope of the EBOV and nucleocapsid. The EBOV genome contains seven structural proteins encoding genes. These proteins are as follows, in 3’- 5’ order: nucleoprotein (NP), glycoprotein (GP), matrix protein (VP24), protein VP30, polymerase cofactor (VP35), matrix protein (VP40), and RNA-dependent RNA polymerase (L). Additionally, there is a small and non-structural glycoprotein called sGP, whose complete function is not fully understood. Although not a component of the virion, this small glycoprotein is released abundantly from infected cells. It may prevent an effective immune response by evading the immune system. Furthermore, the viral proteins VP24 and VP35 serve as significant virulence factors as they function as antagonists to type I interferon (IFN), impeding its function (Feldman et al., 2013; Mateo et al., 2011).

The initial stage of viral replication involves attaching to the host cell membrane and entering inside the cell. Although the exact mechanisms are not completely understood, it has been proved that GP spikes play a role in facilitating the entry of virions into the host cells, potentially through processes similar to macropinocytosis (Aleksandrowicz et al., 2011). Additionally, VP30 has been identified as crucial for the reinitiation of transcription of subsequent genes during viral replication (Biedenkopf et al., 2013). Due to its essential role in these processes, VP30 presents an intriguing target for potential antiviral therapies (Ascenzi et al., 2008). The assembly of viral nucleocapsid requires the presence of matrix protein VP24, as well as NP and VP35. Silencing the expression of matrix protein (VP24) prevents the discharge of viruses, and it also leads to diminished transcription and translation of VP30 in VP24-deficient viral particles (Hoenen et al., 2006). Additionally, the VP40 matrix protein, which is expressed in large amounts, plays a significant role in the formation of novel virus particles. It is closely associated with the endosomal pathway and the process of budding of virus from the host cell (Stahelin 2014).

In regions with tropical climates, where several febrile illnesses can present similar symptoms to Ebola virus disease (EVD), it is very important to consider testing for or providing empirical treatment for parasitic diseases (such as *Plasmodium* spp.), viral diseases, and bacterial diseases (such as *Salmonella typhi*) (Vernet, 2017; Carroll, 2017). While assays of polymerase chain reaction (PCR) are accurate, various factors like cost, processing time (including transportation time of sample), its availability, and the required level of experience of operators have contributed to delays in obtaining rapid diagnostic results during the 2013-2016 EVD outbreak in Western Africa. In response, the WHO called for the development of "rapid, sensitive, safe, and simple EBOV diagnostic tests" in November 2014 (Nouvellet 2015). Mathematical modeling was employed to assess the impact on the case fatality rate (CFR) of EVD and the transmission dynamics of EBOV by incorporating different diagnostic strategies, including the introduction of the novel rapid diagnostic tests (RDTs) (Wannier 2019). Given the current COVID-19 pandemic produced by the SARS-CoV-2 virus, it is becoming increasingly important to conduct research on the specific characteristics of the EBOV that may heighten its potential for causing a worldwide pandemic in the future.

Currently, two licensed vaccines, a two-dose combination of Zabdeno (Ad26.ZEBOV) and Mvabea (MVA-BN-Filo), and Ervebo (rVSV-ZEBOV) are being utilized for EBOV (Tomori and Kolawole 2021). Codon optimization of DNA is a valuable technology used for enhancing the production of foreign proteins. This technique involves modifying the nucleotide sequence of a gene of interest to improve its translational efficiency in a different species, such as transforming a plant sequence into a human sequence or a human sequence into bacterial or yeast sequences (Graf et al., 2009). Codon usage variation is considered an important factor that influences protein expression levels (Gustafsson et al., 2004). The presence of odd codons can slow down the translation process and lead to errors, significantly impacting the efficiency of production processes by recombinant microbe-based (Gustafsson et al., 2004; Heinzelman et al., 2009; Ikemura, 1981).

The objective of our study was to optimize the codon level of all seven genes of EBOV in *Escherichia coli* (*E. coli*) using computational methods. This optimization aimed to achieve higher expression levels of the desired proteins while maintaining their antigenicity and functional activity identical to their native counterparts. By optimizing the DNA codons of the studied genes, we can increase the expression of these proteins, ensuring their efficient and increased production for purposes such as immunodiagnostics and immunotherapy, without introducing changes to their amino acid sequences.

## 2. Materials and method

### 2.1 Collection of sequence

The nucleotide sequences of various strains of the Ebola virus were retrieved from the NCBI-GenBank. The sequences were identified by their accession numbers: KY786027.1, MG572235.1, FJ968794.1, MH121167.1, KY008770.1, and MT742157.1 (http://www.ncbi.nlm.nih.gov).

### 2.2 Analysis and optimization of codons

The online PHP application called Optimizer (http://genomes.urv.es/OPTIMIZER/) (Puigbo et al., 2007) proves to be a valuable tool for predicting and enhancing the expression of a gene in a heterologous gene expression host. Using *Escherichia coli* str. K-12 substr. MG1655 as a reference host, Optimizer was used to optimize and determine the codon adaptation index (CAI), G+C content, and A+T content of the obtained DNA sequences. This strain of *E. coli* is frequently used for heterologous gene expression. CAI values were also determined for each gene across the six distinct strains.

### 2.3 Statistical analysis

The statistical analysis of the genes was conducted using Microsoft Excel 2021 software. Mean, standard deviation, and range calculations were performed on the gene data. The data were grouped in a table, and a graph was generated to illustrate the comparison between the CAI values of wild-type and optimized gene sequences for various EBOV variants. The Mann-Whitney test was used to compare the CAI, GC content, and AT content of all seven genes. For statistical significance, a two-tailed probability p < 0.05 was used.

### 2.4 Nucleotide sequence alignment

The nucleotide sequences of all seven genes from strains KY786027.1, MG572235.1, FJ968794.1, MH121167.1, KY008770.1, and MT742157.1 were aligned using Optimizer. The alignment was performed between the wild-type sequences and the corresponding optimized sequences for each gene.

## 3. Results

The present study aimed to address this need by employing DNA codon optimization to produce sufficient quantities of proteins in the desired host. In our study, the CAI of the optimized DNA sequences was found to be greater than that of the wild-type sequences. The CAI, GC% (percentage of guanine and cytosine) and AT% (percentage of adenine and thymine) for the wild-type nucleoprotein gene varied among the six different strains, ranging from 0.3 to 0.319, 45.4 to 46.9, and 53.1 to 54.6, respectively. The average values (± standard deviation) for CAI, GC%, and AT% were 0.319 (±0.007), 46.16 (±0.668), and 53.83 (±0.668), respectively. On the other hand, the frequencies of GC% and AT% in the optimized DNA ranged from 55.1 to 55.91 and 44.1 to 46.2, respectively, with average values (± standard deviation) of 55.01 (±0.685) and 44.98 (±0.685). Notably, the CAI for the optimized DNA was 1 for all six strains, indicating an enhanced translational efficiency of the optimized sequences.

Upon comparing the average values of the CAI, AT content, and GC content of the nucleoprotein gene across all six strains, notable differences were observed between the optimized DNA and the wild-type sequences. The optimized DNA exhibited significantly higher values for GC and CAI content, with increases of 1.2 times (19.17%) and 3.14 times (213.5%), respectively, compared to the mean values of the wild-type sequences. Conversely, the optimized DNA displayed a reduction of 16.4% in the mean AT content compared to the wild-type sequences (Table 1). To visually represent the variations in CAI values among the studied strains, a graph was generated, plotting CAI values along the x-axis and the number of strains along the y-axis (Fig. 1A). Furthermore, an alignment was performed on the nucleocapsid gene sequences of both the codon-optimized sequences and wild-type sequences, as illustrated in Fig.1B. Importantly, it should be emphasized that no alteration was seen in the amino acid sequence of the nucleoprotein by the codon optimization process. Its goal was to improve translational efficiency in order to improve the immunogenicity of epitope-based vaccinations. As a result, altering the codon bias of gene sequences provides a possible technique for influencing gene expression.

**Table 1:** The expression level of NP gene of the Ebola virus in *E. coli* of wild-type and codon-optimized sequences.

| Ebola virus strains<br>(GenBank Accession No.) | Wild-Type DNA |  |  | Optimized DNA |  |  |
| --- | --- | --- | --- | --- | --- | --- |
|  | CAI | GC% | AT% | CAI | GC% | AT% |
| KY786027.1 | 0.319 | 46.9 | 53.1 | 1 | 55.2 | 44.8 |
| MG572235.1 | 0.311 | 46.9 | 53.1 | 1 | 55.2 | 44.8 |
| FJ968794.1 | 0.312 | 46.4 | 53.6 | 1 | 54.9 | 45.1 |
| MH121167.1 | 0.305 | 45.4 | 54.6 | 1 | 53.8 | 46.2 |
| KY008770.1 | 0.3 | 45.5 | 54.5 | 1 | 55.9 | 44.1 |
| MT742157.1 | 0.311 | 45.9 | 54.1 | 1 | 55.1 | 44.9 |
| **N** | 6 | 6 | 6 | 6 | 6 | 6 |
| **Min.** | 0.3 | 45.4 | 53.1 | 1 | 55.1 | 44.1 |
| **Max.** | 0.319 | 46.9 | 54.6 | 1 | 55.9 | 46.2 |
| **Mean±SD** | 0.319±0.007 | 46.16±0.668 | 53.83±0.668 | 1.00±0.00 | 55.01±0.685 | 44.98±0.685 |

**Fig. 1A:**
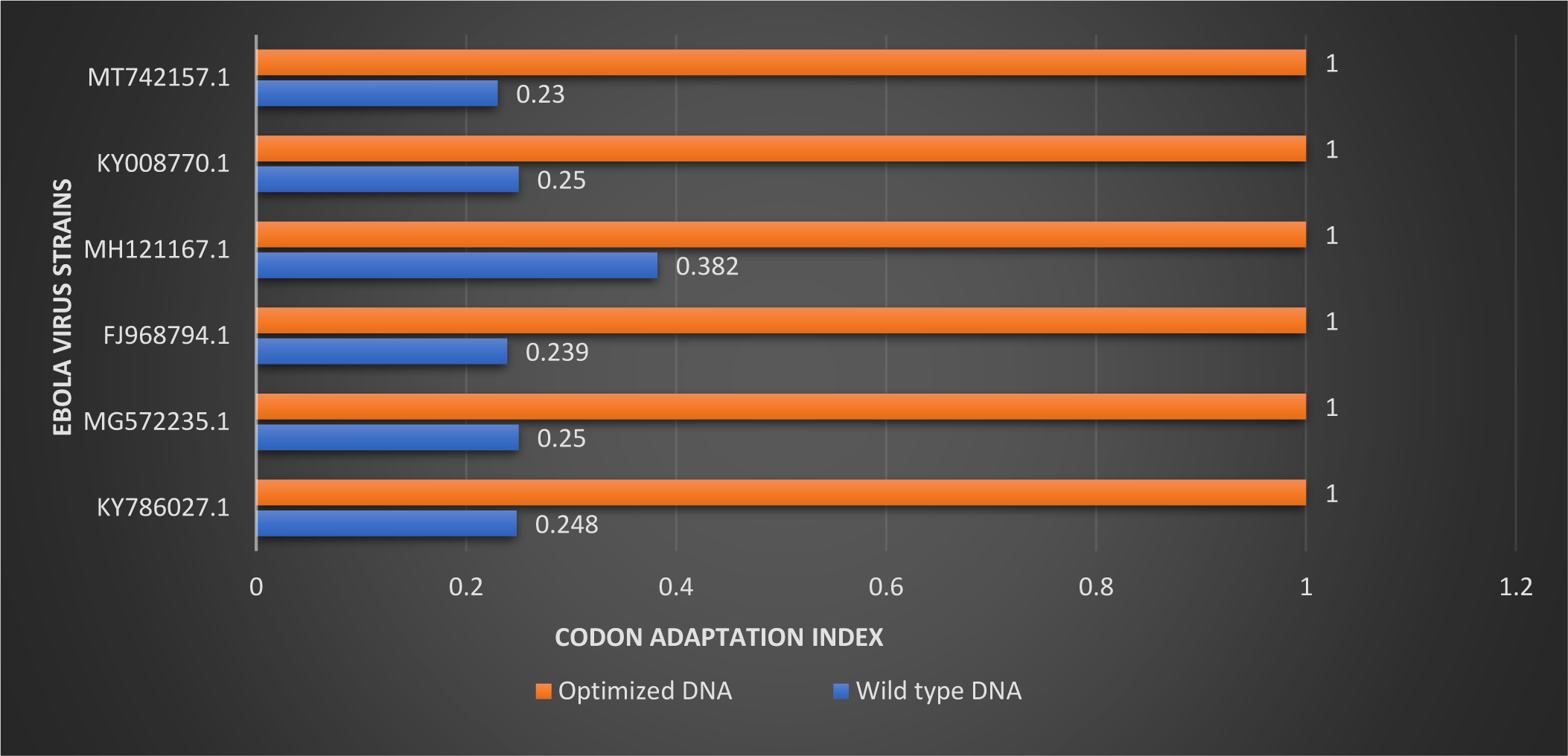
Graph showing comparison between the wild-type and optimized DNA for nucleocapsid (N) gene.

**Fig. 1B:**
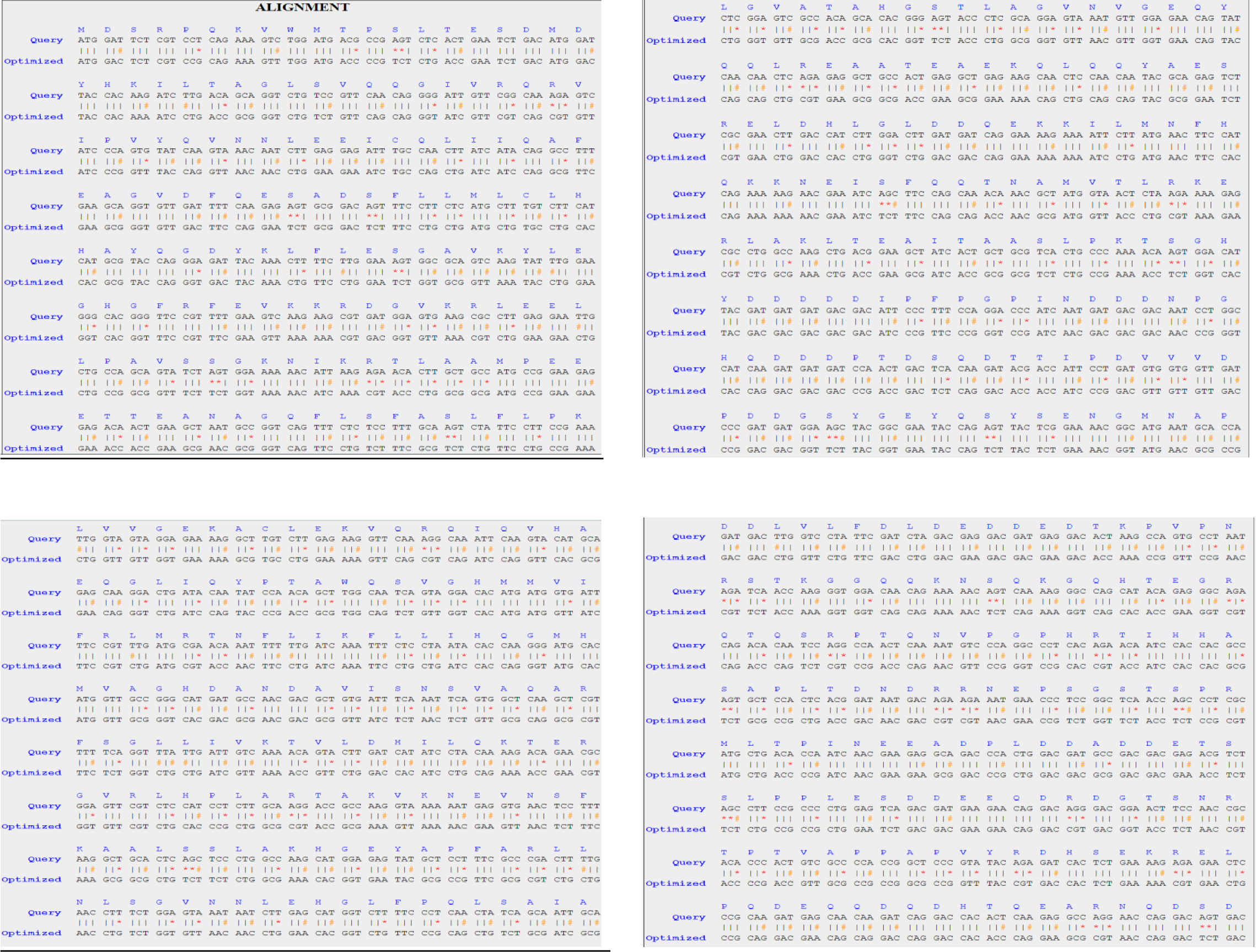
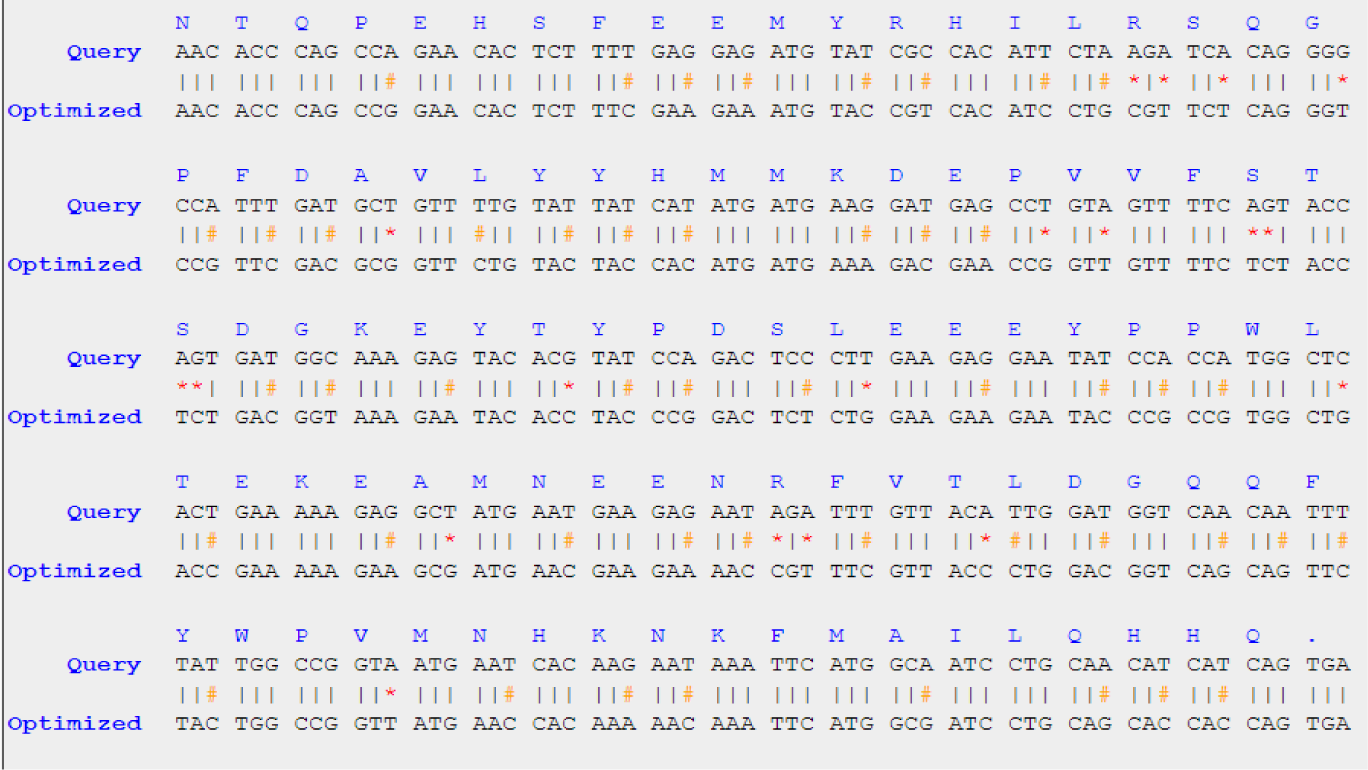
The sequence alignment of wild-type (Query) and codon optimized DNA for nucleocapsid (NP) gene of Ebola virus for accession number MG572235.1. *The specific-coloured symbols stand for (*): Transversion change, (#): Transition change and (|): Unchanged nucleotide

In the case of the VP35 gene, the CAI, AT content, and GC content in the wild-type sequences of the six different strains ranged from 0.269 to 0.309, 41 to 46.8, and 53.2 to 59, respectively. The average values (±SD) were 0.29 (±0.013) for CAI, 44.78 (±1.996) for GC content, and 55.21 (±1.996) for AT content (Table 2). Upon optimization, the frequencies of GC and AT content in the respective DNA sequences ranged from 51.4 to 56.4 and 43.6 to 48.6, with average values (±SD) of 54.86 (±1.949) and 45.13 (±1.949), respectively. The CAI for the optimized DNA was 1 for all six strains.

**Table 2:** The expression level of VP35 gene of the Ebola virus in *E. coli* of wild type and codon-optimized sequences.

| Ebola virus strains<br>(GenBank Accession No.) | Wild-Type DNA |  |  | Optimized DNA |  |  |
| --- | --- | --- | --- | --- | --- | --- |
|  | CAI | GC% | AT% | CAI | GC% | AT% |
| KY786027.1 | 0.299 | 44.8 | 55.2 | 1 | 56.1 | 43.9 |
| MG572235.1 | 0.3 | 45.5 | 54.5 | 1 | 56.4 | 43.6 |
| FJ968794.1 | 0.289 | 44.8 | 55.2 | 1 | 53.8 | 46.2 |
| MH121167.1 | 0.297 | 46.8 | 53.2 | 1 | 56.2 | 43.8 |
| KY008770.1 | 0.269 | 41 | 59 | 1 | 51.4 | 48.6 |

When the mean values of CAI, GC content, and AT content for the VP35 gene in all six strains were compared, the optimized DNA had considerably higher values. The optimized DNA’s mean CAI and GC content were found to be 3.44 (244.8%) and 1.22 (22.5%) times greater, respectively, than the wild-type sequences’ corresponding mean values. The mean AT content in the optimized DNA, on the other hand, was reduced by 18.25% when compared to the wild-type sequences (Table 2). A graph was created to show these data (Fig. 2A). Both the wild-type and codon-optimized VP35 gene sequences were aligned, as shown in Fig. 2B.

**Fig. 2A:**
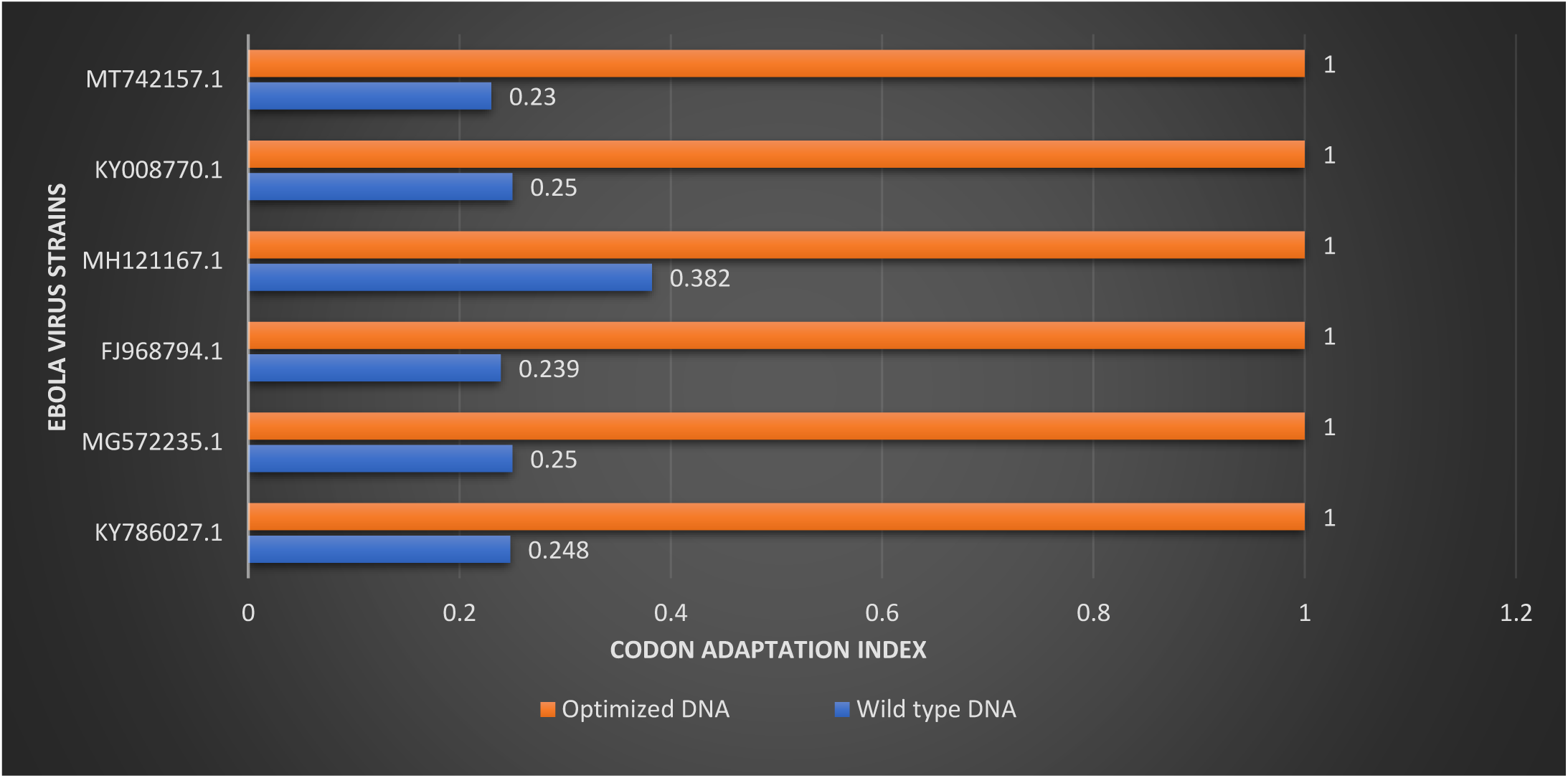
Graph showing comparison between the wild-type and optimized DNA for VP35 gene.

**Fig. 2B:**
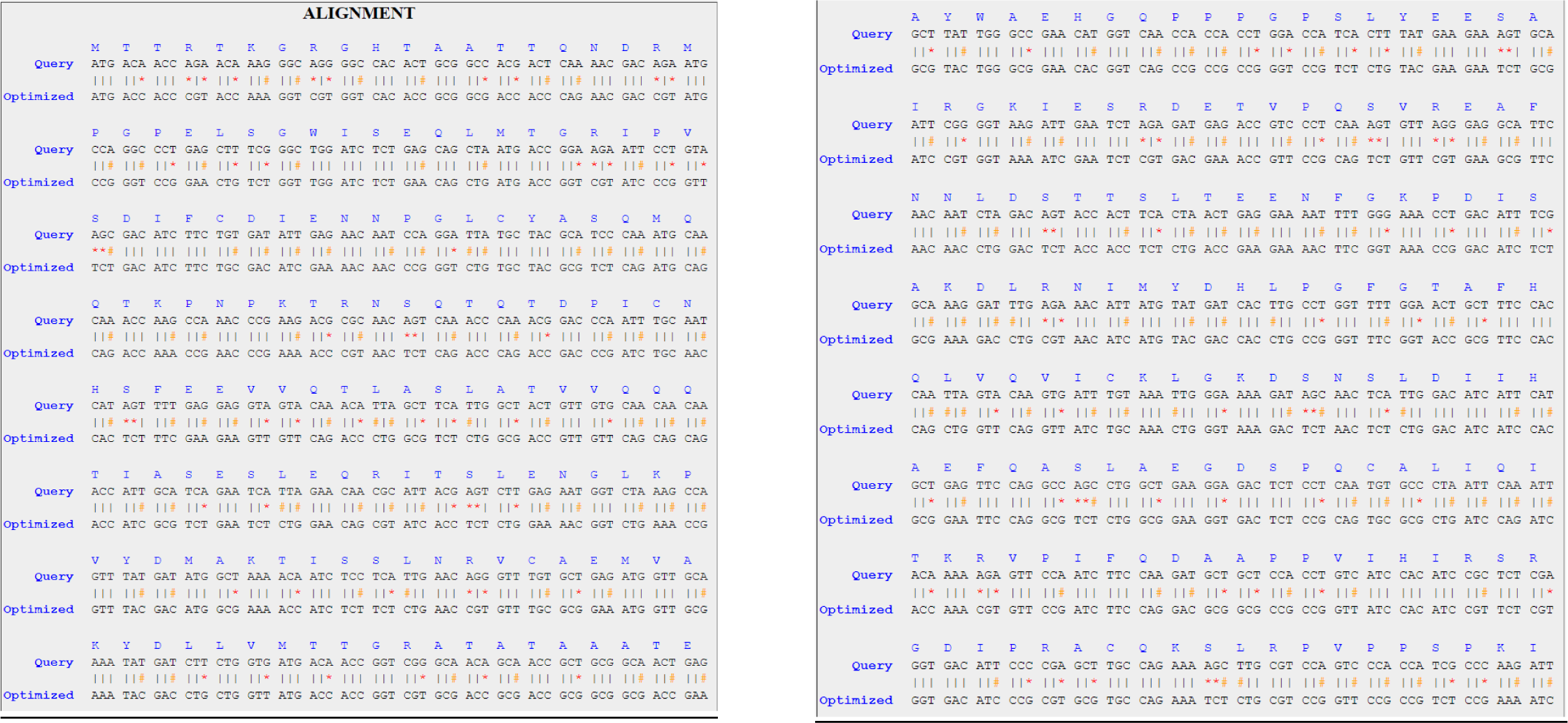
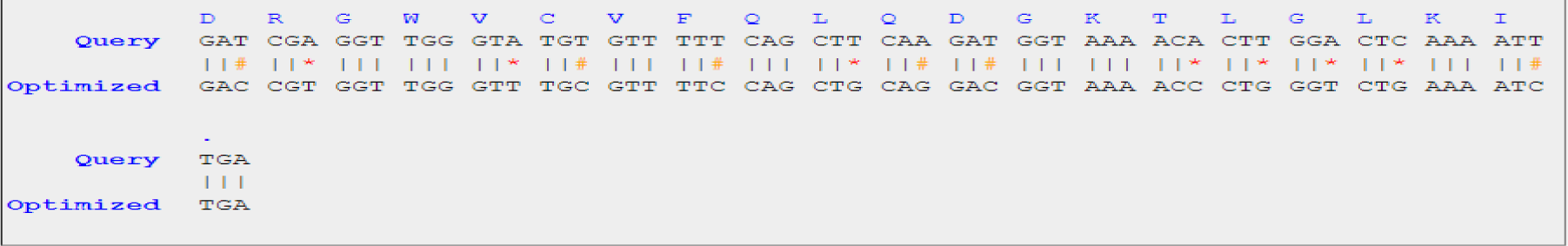
The sequence alignment of wild-type (Query) and codon optimized DNA for VP35 gene of Ebola virus for accession number MG572235.1. *The specific-coloured symbols stand for (*): Transversion change, (#): Transition change and (|): Unchanged nucleotide

For the VP40 gene, the CAI, GC content, and AT content in the wild-type sequences of the six different strains ranged from 0.246 to 0.278, 45.7 to 48.6, and 51.4 to 54.3, respectively. The average values (±SD) were 0.26 (±0.014) for CAI, 47.13 (±1.169) for GC content, and 52.86 (±1.169) for AT content (Table 3). After codon optimization, the frequencies of GC and AT content in the respective DNA sequences ranged from 56.6 to 57.7 and 42.3 to 43.4, with average values (±SD) of 57.12 (±0.36) and 42.88 (±0.360), respectively. The CAI for the optimized DNA was 1 for all six strains.

**Table 3:**
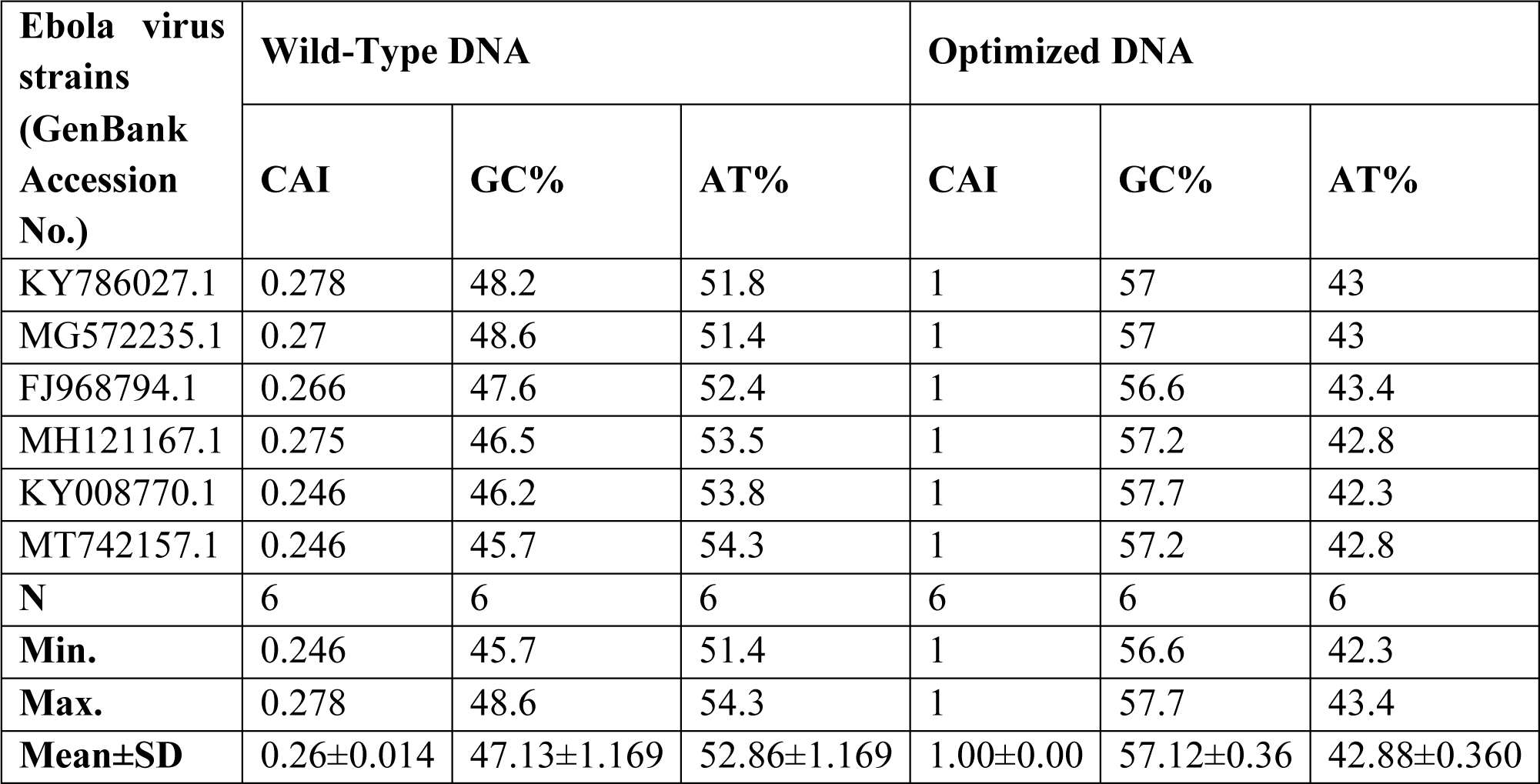
The expression level of VP40 gene of Ebola virus in *E. coli* of wild-type and codon-optimized sequences.

Upon comparing the mean values of CAI, GC content, and AT content for the VP40 gene in all six strains, it was observed that the values of the optimized DNA were significantly higher. The optimized DNA’s mean CAI and GC content were found to be 3.84 (284.6%) and 1.2 (21.2%) times greater, respectively, than the wild-type sequences’ corresponding mean values. The mean AT content in the optimized DNA, on the other hand, was reduced by 18.88% when compared to the wild-type sequences (Table 3). To visualize these findings, a graph was created (Fig. 3A). Both the wild-type and codon-optimized VP40 gene sequences were aligned, as shown in Fig. 3B. It is worth mentioning that codon optimization did not result in any changes to the VP40 gene’s amino acid sequence.

**Fig. 3A:**
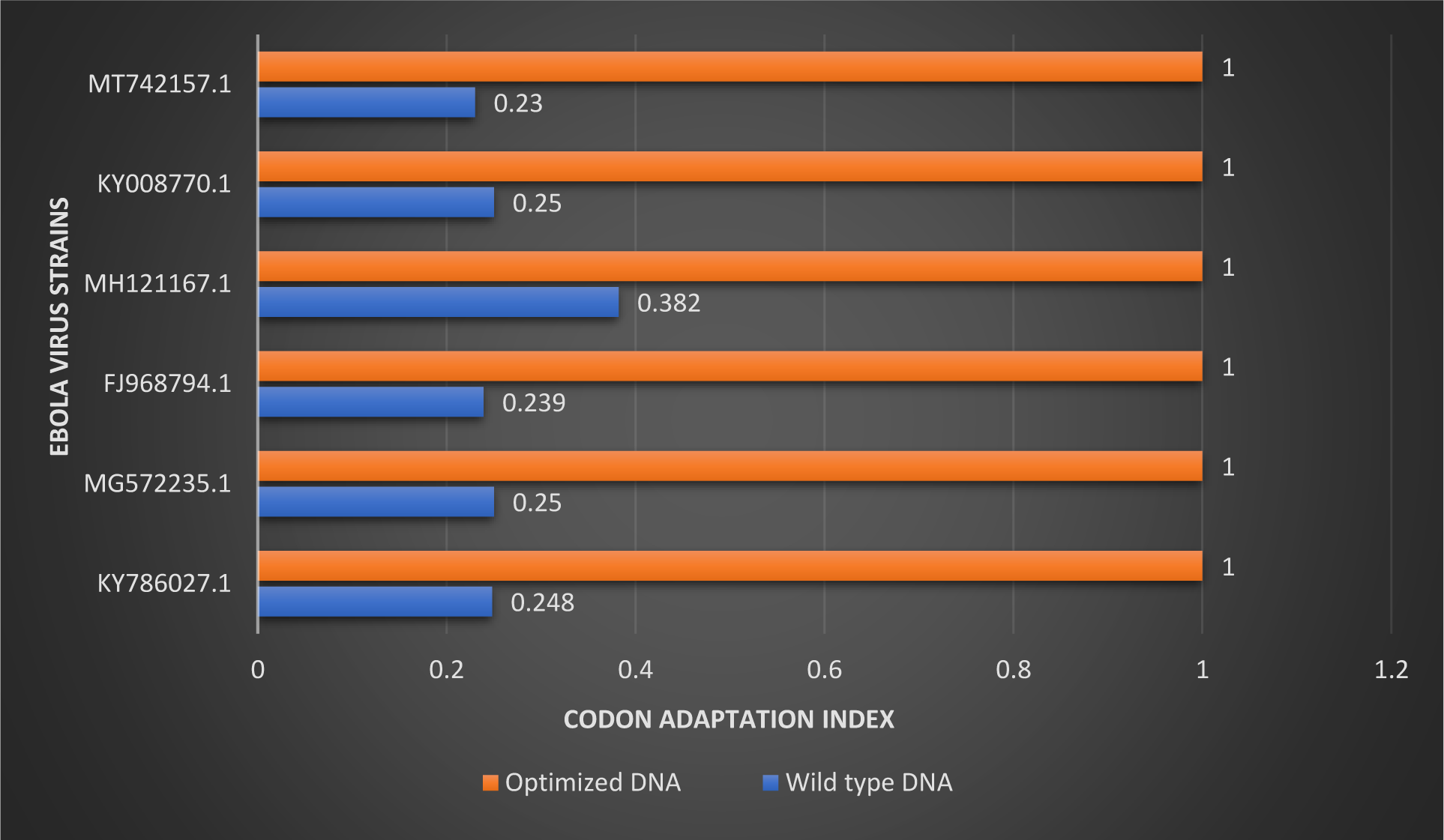
Graph showing comparison between the wild-type and optimized DNA for VP40 gene.

**Fig. 3B:**
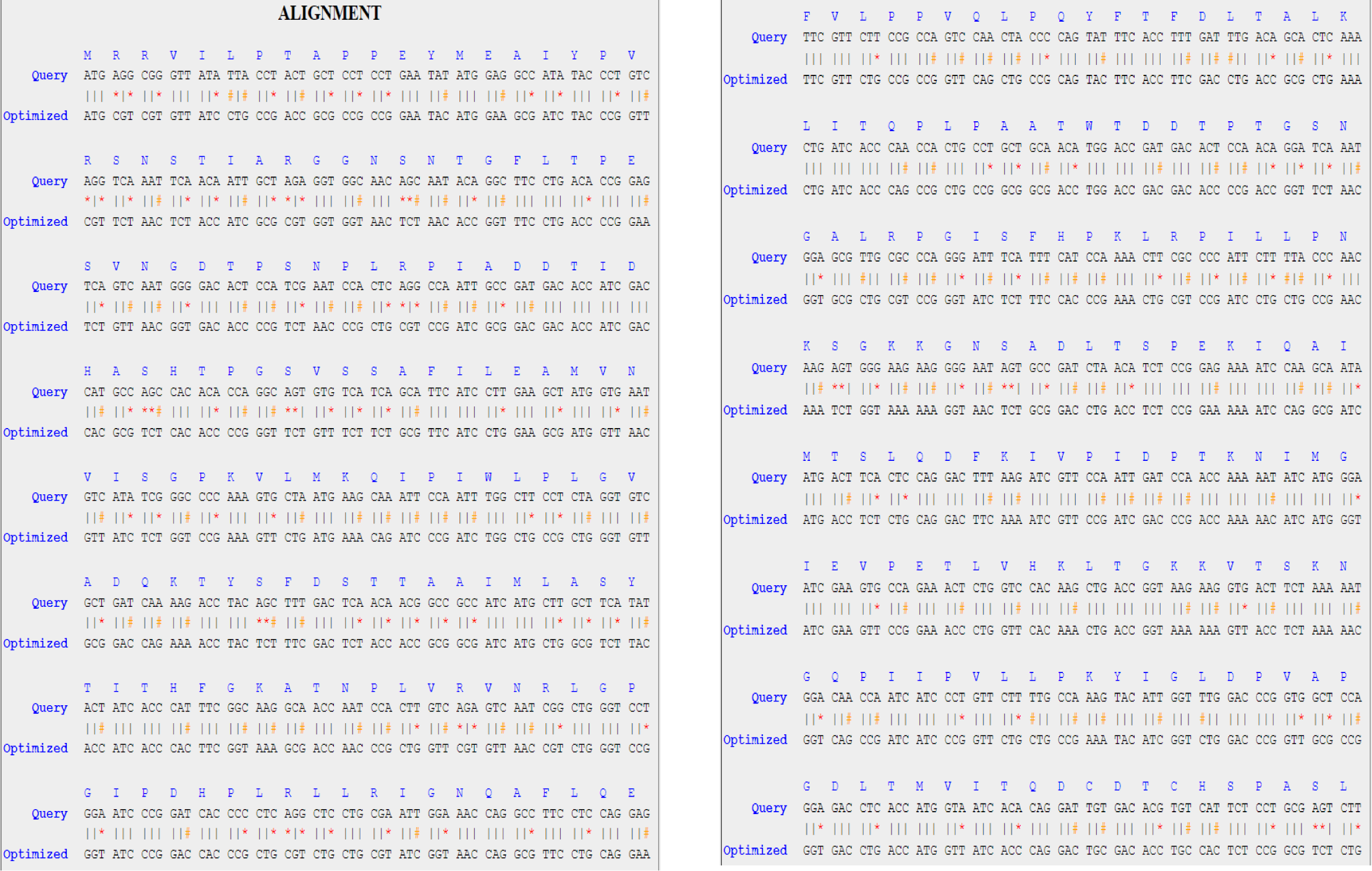
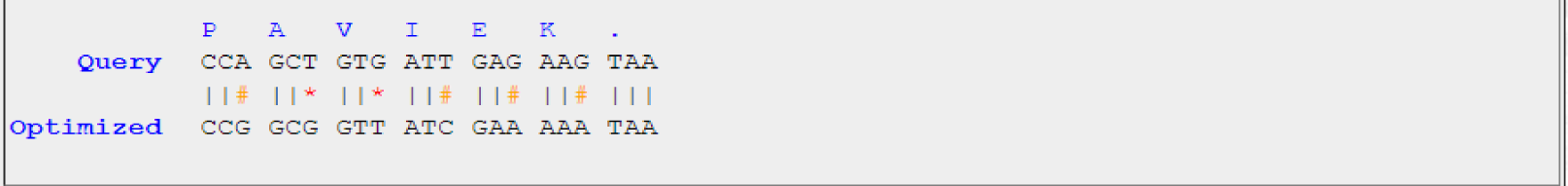
The sequence alignment of wild-type (Query) and codon-optimized DNA for VP40 gene of Ebola virus for accession number MG572235.1. *The specific-coloured symbols stand for (*): Transversion change, (#): Transition change and (|): Unchanged nucleotide

The wild-type GP gene exhibited a range of 0.26 to 0.295 for CAI, 46 to 46.8 for GC content, and 53.2 to 54 for AT content across the six different strains. The average values (±SD) were 0.28 (±0.012) for CAI, 46.48 (±0.279) for GC content, and 53.52 (±0.279) for AT content (Table 4). After codon optimization, the frequencies of GC and AT content in the respective DNA sequences ranged from 53 to 55.4 and 44.6 to 47, with average values (±SD) of 54.18 (±0.813) and 45.82 (±0.813), respectively. The CAI for the optimized DNA was 1 for all six strains.

**Table 4:** The expression level of GP gene of Ebola virus in *E. coli* of wild-type and codon optimized sequences.

| Ebola virus strains<br>(GenBank Accession No.) | Wild-Type DNA |  |  | Optimized DNA |  |  |
| --- | --- | --- | --- | --- | --- | --- |
|  | CAI | GC% | AT% | CAI | GC% | AT% |
| KY786027.1 (sGP) | 0.275 | 46.8 | 53.2 | 1 | 53 | 47 |
| MG572235.1 | 0.276 | 46 | 54 | 1 | 54.7 | 45.3 |
| FJ968794.1 | 0.286 | 46.7 | 53.3 | 1 | 55.4 | 44.6 |
| MH121167.1 | 0.26 | 46.5 | 53.5 | 1 | 53.8 | 46.2 |
| KY008770.1 | 0.288 | 46.5 | 53.5 | 1 | 54.1 | 45.9 |
| MT742157.1 | 0.295 | 46.4 | 53.6 | 1 | 54.1 | 45.9 |

Upon comparing the mean values of CAI, GC content, and AT content for the glycoprotein gene in all six strains, it was observed that the values of the optimized DNA were significantly higher. The optimized DNA’s mean CAI and GC content were found to be 3.57 (257.14%) and 1.16 (16.56%) times higher, respectively, than the wild-type sequences’ corresponding mean values. The mean AT content in the optimized DNA, on the other hand, was reduced by 14.38% when compared to the wild-type sequences (Table 4). To visualize these findings, a graph was created (Fig. 4A). As demonstrated in Fig. 4B, the glycoprotein gene sequences of both the wild-type and codon-optimized sequences were aligned. The goal of codon optimization is to improve translational efficiency and thereby boost the immunogenicity of epitope-based vaccinations. Modifying the codon bias of gene sequences thus bears potential as a method for controlling gene expression.

**Fig. 4A:**
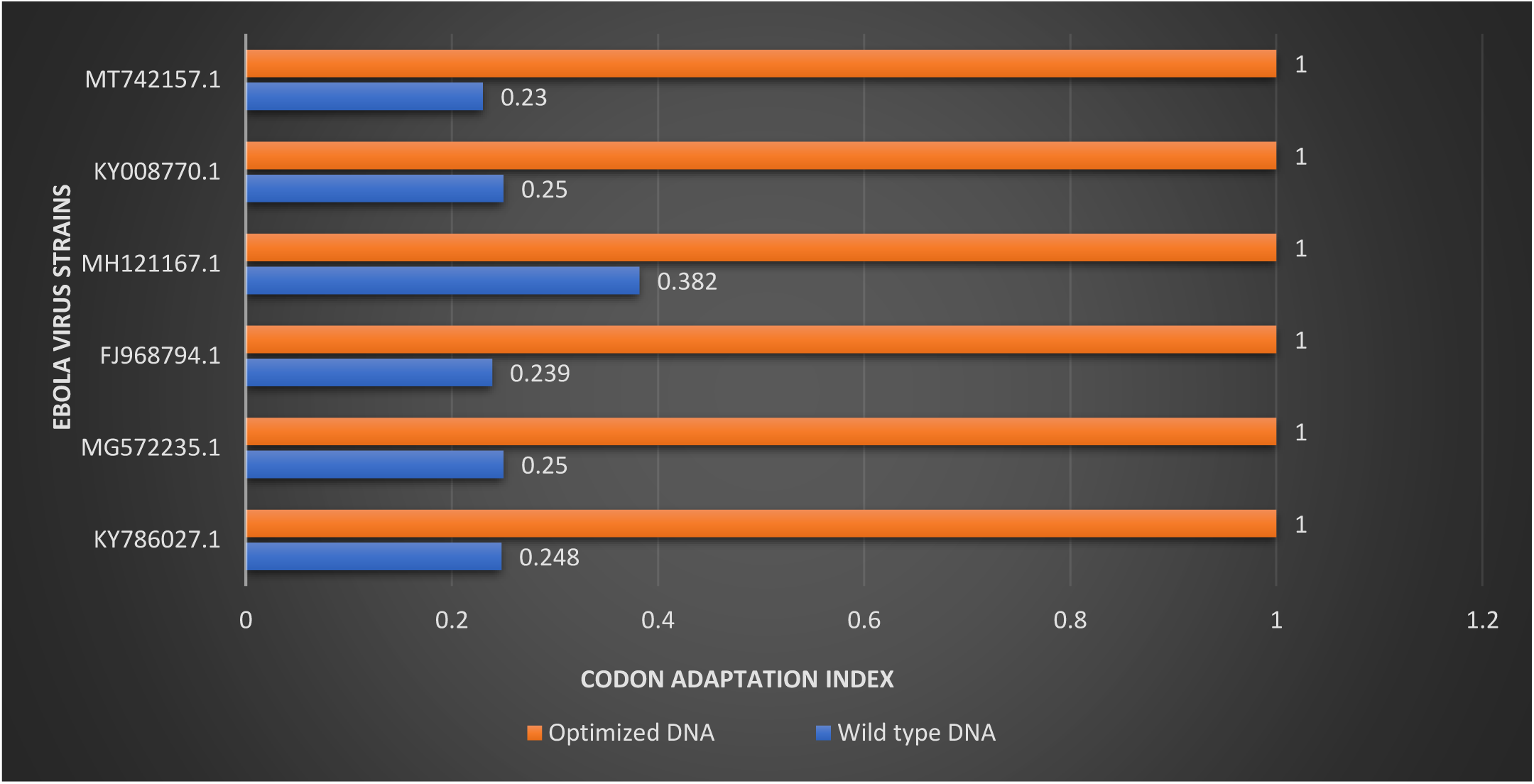
Graph showing comparison between the wild-type and optimized DNA for glycoprotein (GP) gene.

**Fig. 4B:**
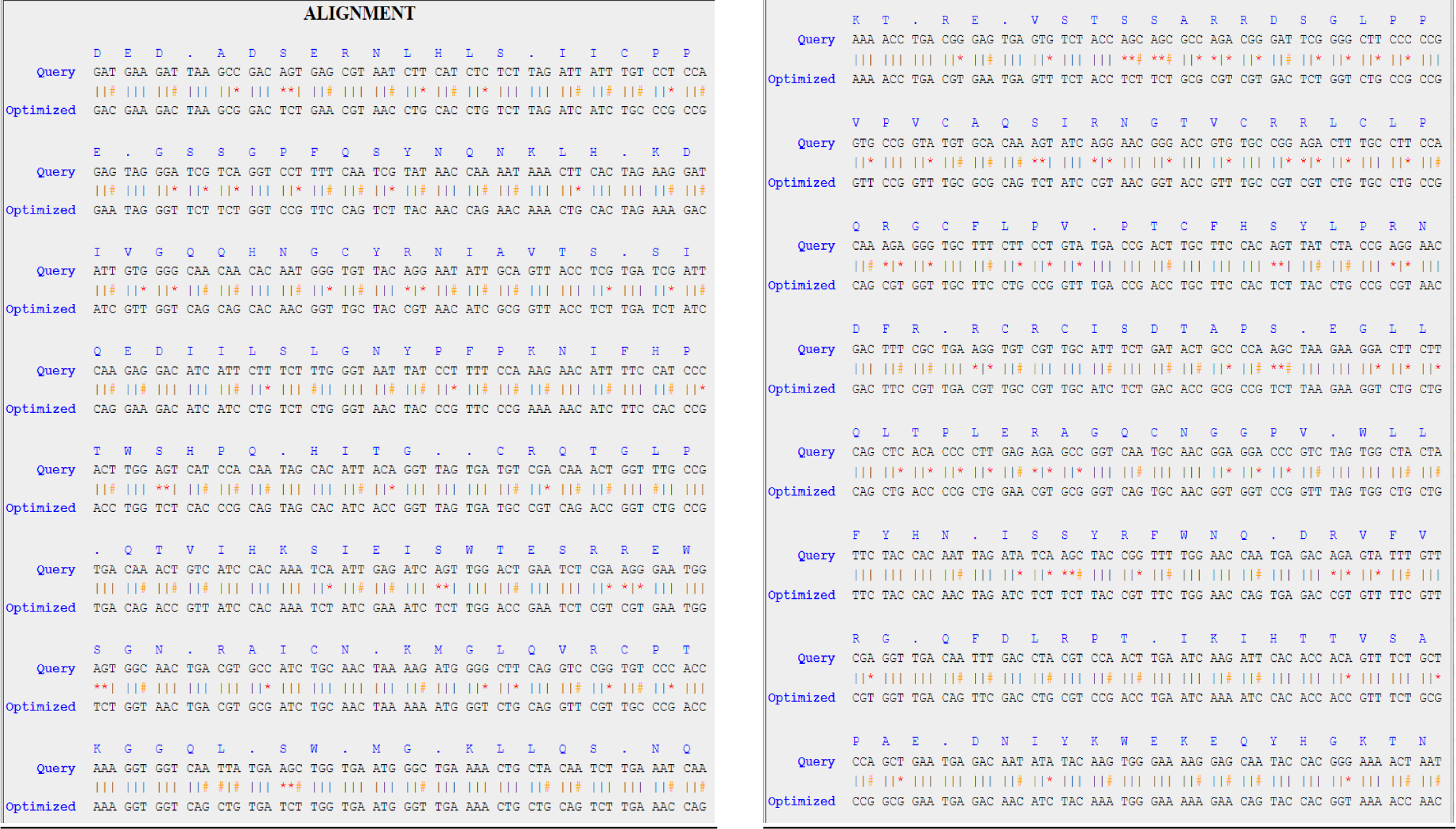
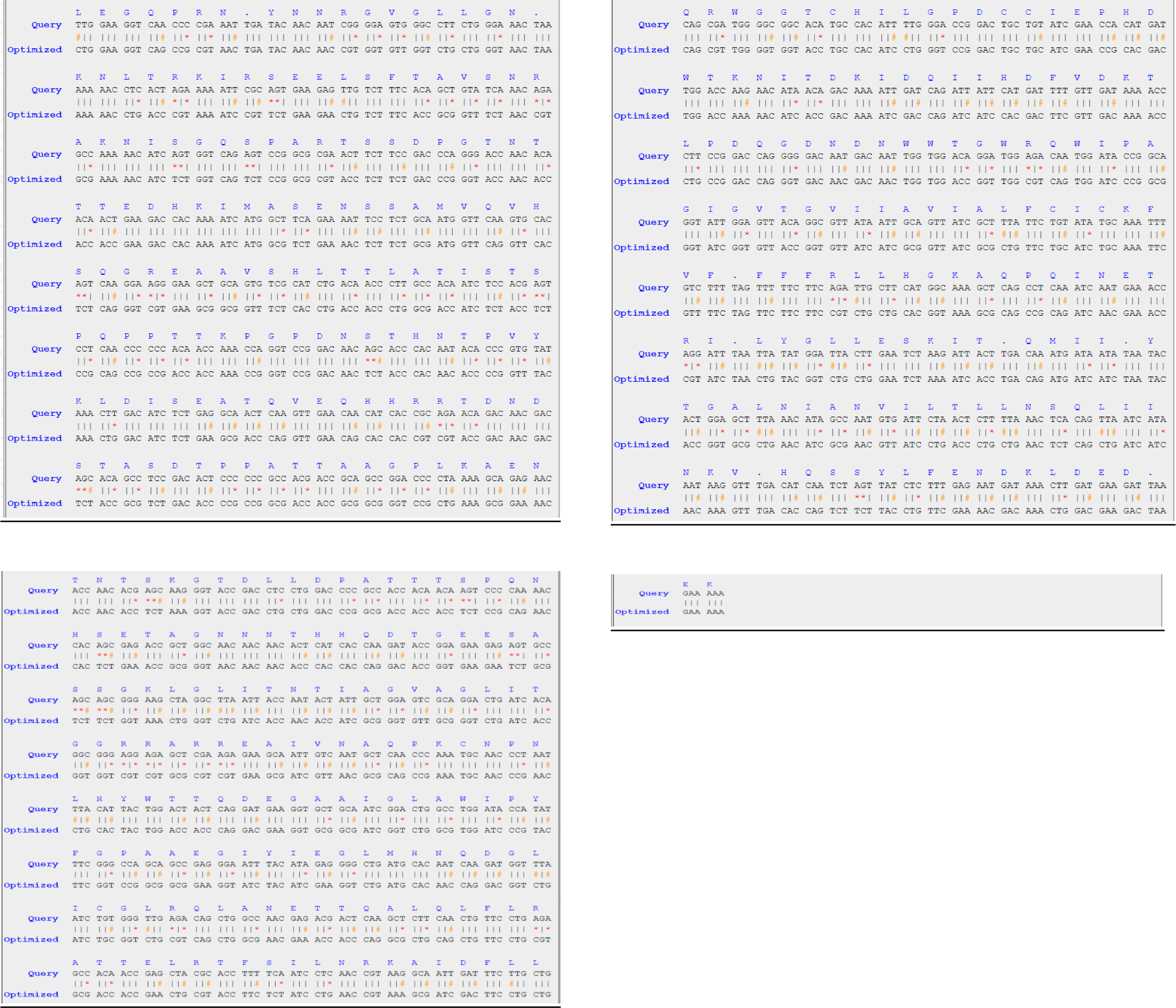
The sequence alignment of wild-type (Query) and codon optimized DNA for glycoprotein (GP) gene of Ebola virus for accession number MG572235.1. *The specific-coloured symbols stand for (*): Transversion change, (#): Transition change and (|): Unchanged nucleotide

The wild-type VP30 gene exhibited a range of 0.225 to 0.261 for CAI, 39.4 to 45.3 for GC content, and 54.7 to 60.6 for AT content across the six different strains. The average values (±SD) were 0.23 (±0.012) for CAI, 43.05 (±2.617) for GC content, and 56.95 (±2.617) for AT content (Table 5). After codon optimization, the frequencies of GC and AT content in the respective DNA sequences ranged from 48.5 to 57.1 and 42.9 to 51.5, with average values (±SD) of 54.26 (±3.322) and 45.73 (±3.322), respectively. The CAI for the optimized DNA was 1 for all six strains.

**Table 5:** The expression level of VP30 gene of Ebola virus in *E. coli* of wild-type and codon-optimized sequences.

| Ebola virus strains<br>(GenBank Accession No.) | Wild-Type DNA |  |  | Optimized DNA |  |  |
| --- | --- | --- | --- | --- | --- | --- |
|  | CAI | GC% | AT% | CAI | GC% | AT% |
| KY786027.1 | 0.234 | 45.3 | 54.7 | 1 | 57.1 | 42.9 |
| MG572235.1 | 0.235 | 44.6 | 55.4 | 1 | 56.4 | 43.6 |
| FJ968794.1 | 0.23 | 45.1 | 54.9 | 1 | 56.1 | 43.9 |
| MH121167.1 | 0.261 | 39.4 | 60.6 | 1 | 48.5 | 51.5 |
| KY008770.1 | 0.231 | 40.1 | 59.9 | 1 | 52.1 | 47.9 |
| MT742157.1 | 0.225 | 43.8 | 56.2 | 1 | 55.4 | 44.6 |
| **N** | 6 | 6 | 6 | 6 | 6 | 6 |
| **Min.** | 0.225 | 39.4 | 54.7 | 1 | 48.5 | 42.9 |
| **Max.** | 0.261 | 45.3 | 60.6 | 1 | 57.1 | 51.5 |
| **Mean±SD** | 0.23±0.012 | 43.05±2.617 | 56.95±2.617 | 1.00±0.00 | 54.26±3.322 | 45.73±3.322 |

When comparing the mean values of CAI, GC content, and AT content for the VP30 gene in all six strains, it was evident that the values of the optimized DNA were significantly higher. The optimized DNA’s mean CAI and GC content were found to be 4.34 (334.8%) and 1.26 (26.04%) times greater, respectively, than the wild-type sequences’ corresponding mean values. The mean AT content in the optimized DNA, on the other hand, was reduced by 19.7% when compared to the wild-type sequences (Table 5). Both the wild-type and codon-optimized VP30 gene sequences were aligned, as shown in Fig. 5B. To visualize these findings, a graph was created (Fig. 5A).

**Fig. 5A:**
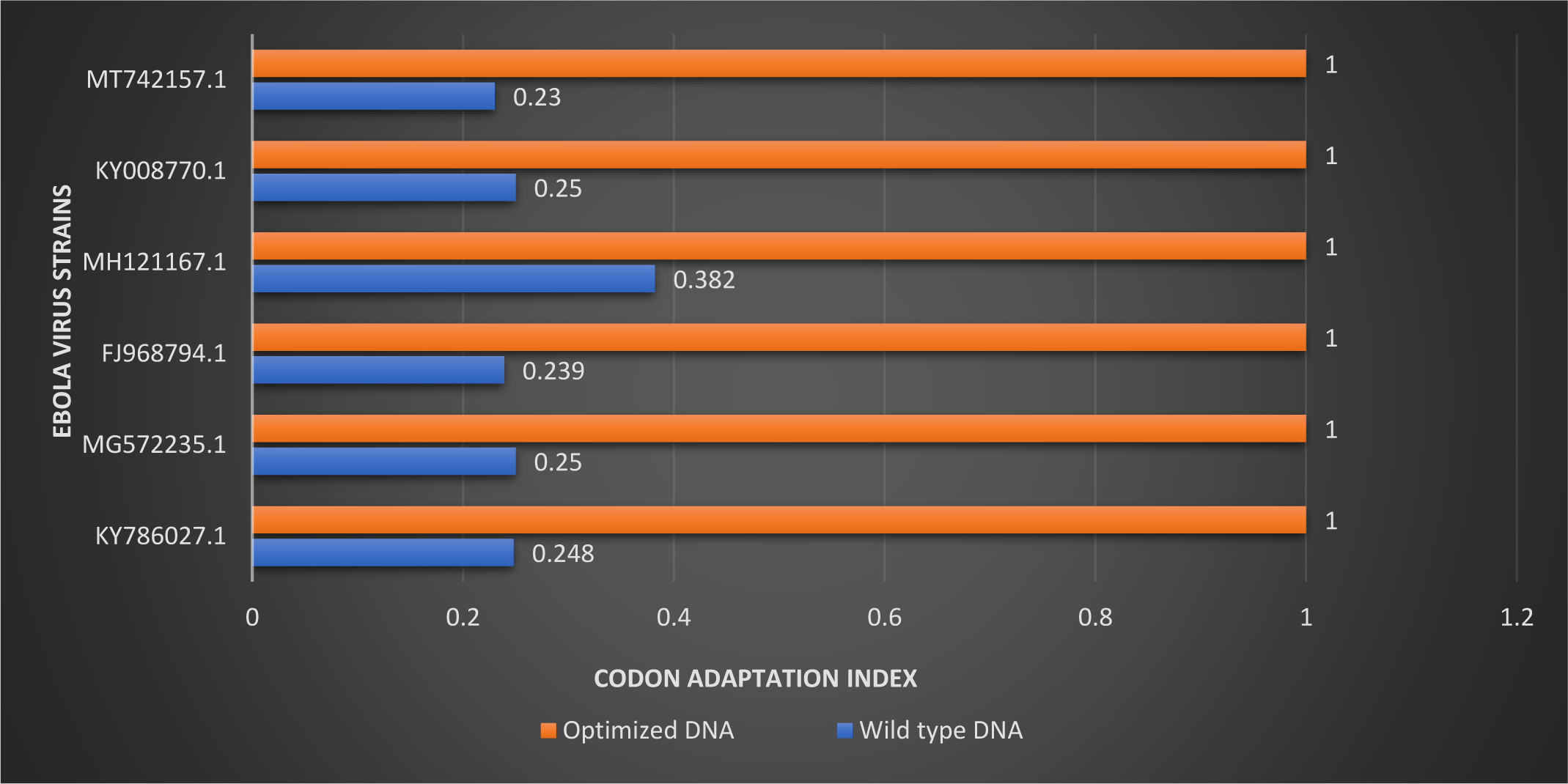
Graph showing comparison between the wild-type and optimized DNA for VP30 gene.

**Fig. 5B:**
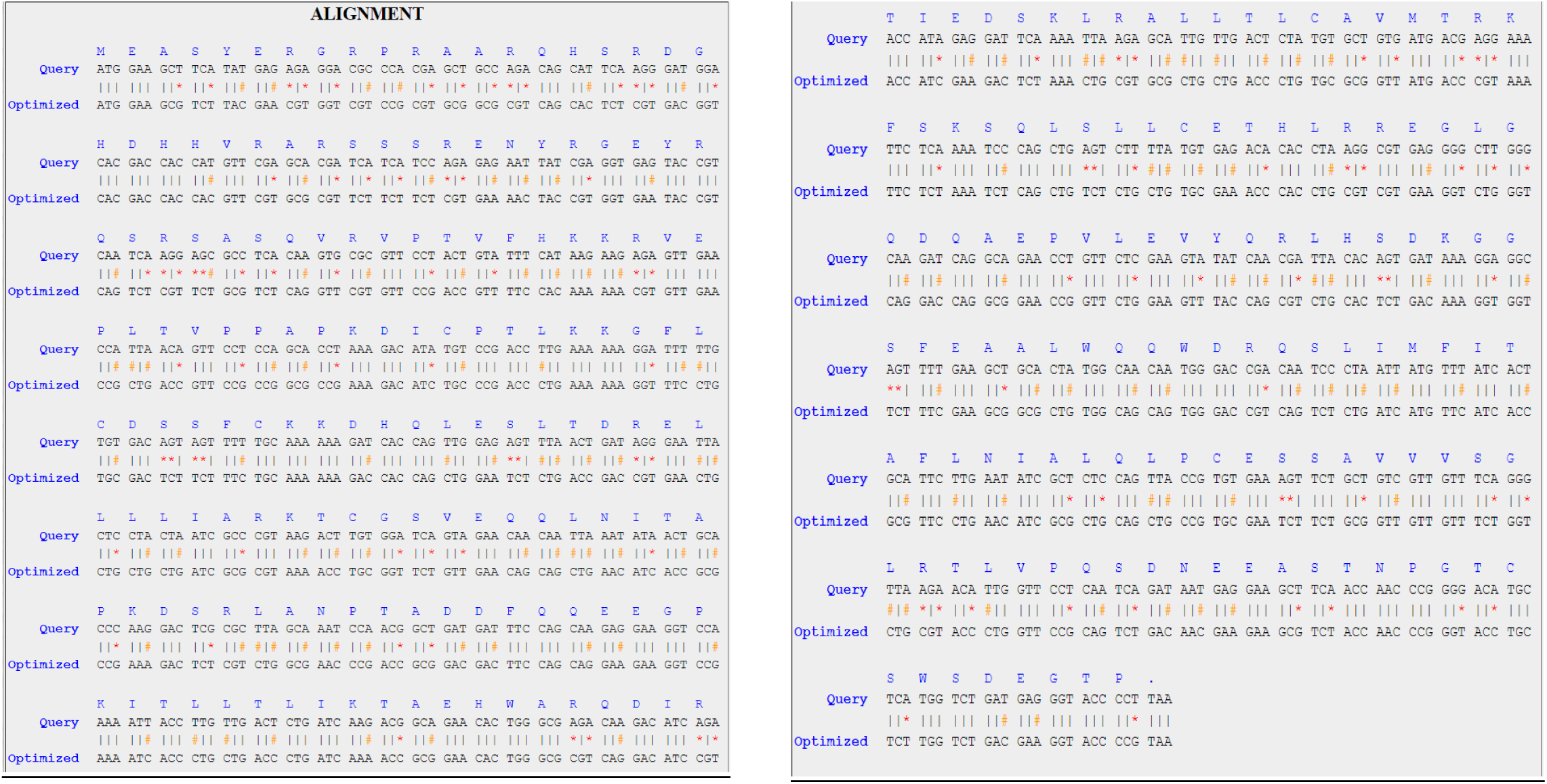
The sequence alignment of wild-type (Query) and codon optimized DNA for VP30 gene of Ebola virus for accession number MG572235.1. *The specific-coloured symbols stand for (*): Transversion change, (#): Transition change and (|): Unchanged nucleotide

The wild-type VP24 gene demonstrated a range of 0.256 to 0.289 for CAI, 41.9 to 44.3 for GC content, and 55.7 to 58.1 for AT content across the six different strains. The average values (±SD) were 0.26 (±0.012) for CAI, 43.06 (±0.973) for GC content, and 56.93 (±0.973) for AT content (Table 6). Following codon optimization, the frequencies of GC and AT content in the respective DNA sequences ranged from 52.4 to 54.2 and 45.8 to 47.6, with average values (±SD) of 53.1 (±0.745) and 46.9 (±0.745), respectively. The CAI for the optimized DNA was 1 for all six strains.

**Table 6:**
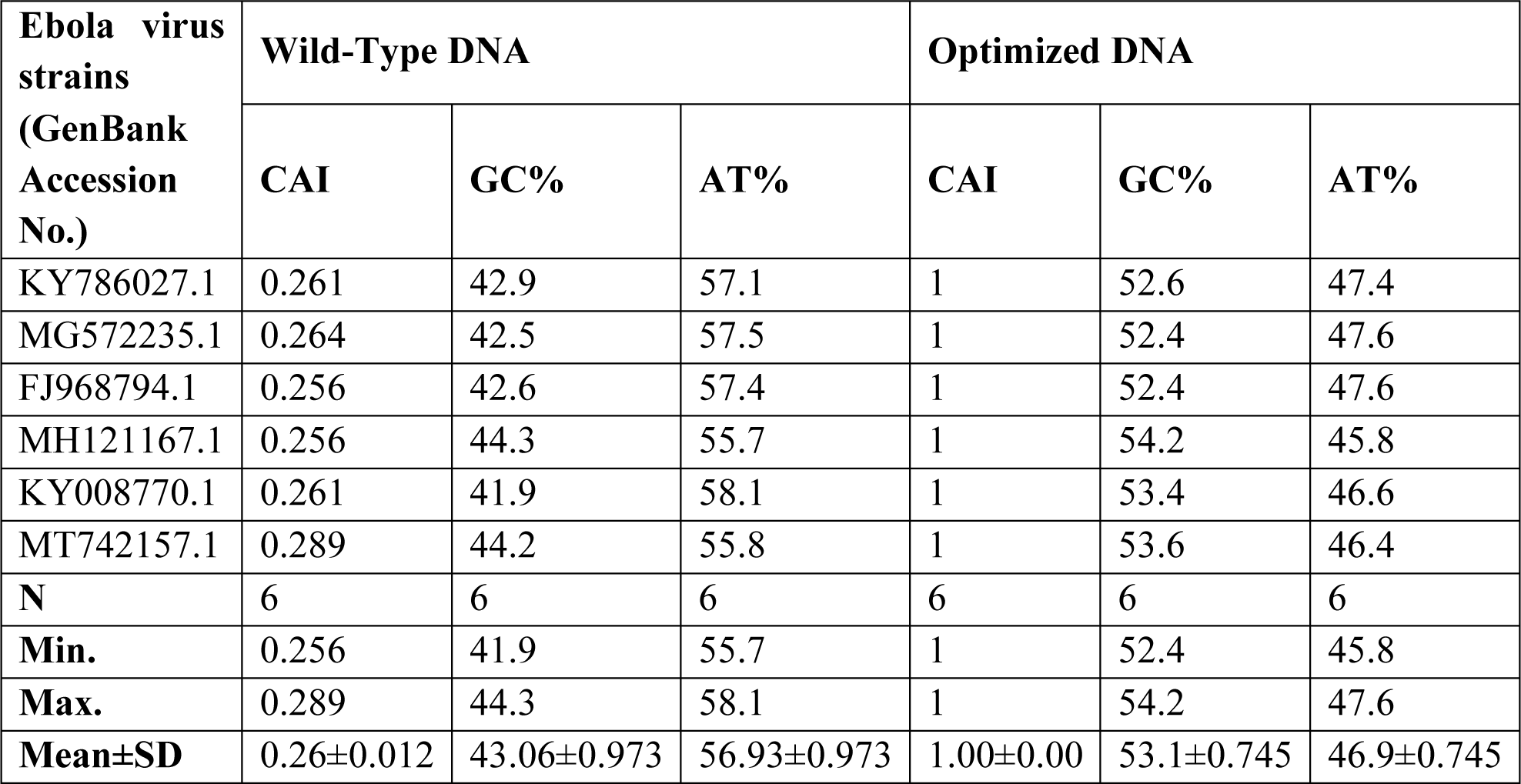
The expression level of VP24 gene of Ebola virus in *E. coli* of wild-type and codon-optimized sequences.

When comparing the mean values of CAI, GC content, and AT content for the VP24 gene in all six strains, it was observed that the values of the optimized DNA were significantly higher. The optimized DNA’s mean CAI and GC content were found to be 3.84 (284.61%) and 1.23 (23.3%) times higher, respectively, than the wild-type sequences’ corresponding mean values. The mean AT content in the optimized DNA, on the other hand, was reduced by 17.61% when compared to the wild-type sequences (Table 6). Both the wild-type and codon-optimized VP24 gene sequences were aligned, as shown in Fig. 6B. To visualize these findings, a graph was created (Fig. 6A). The amino acid sequence of the VP24 gene was not altered as a consequence of codon optimization.

**Fig 6A:**
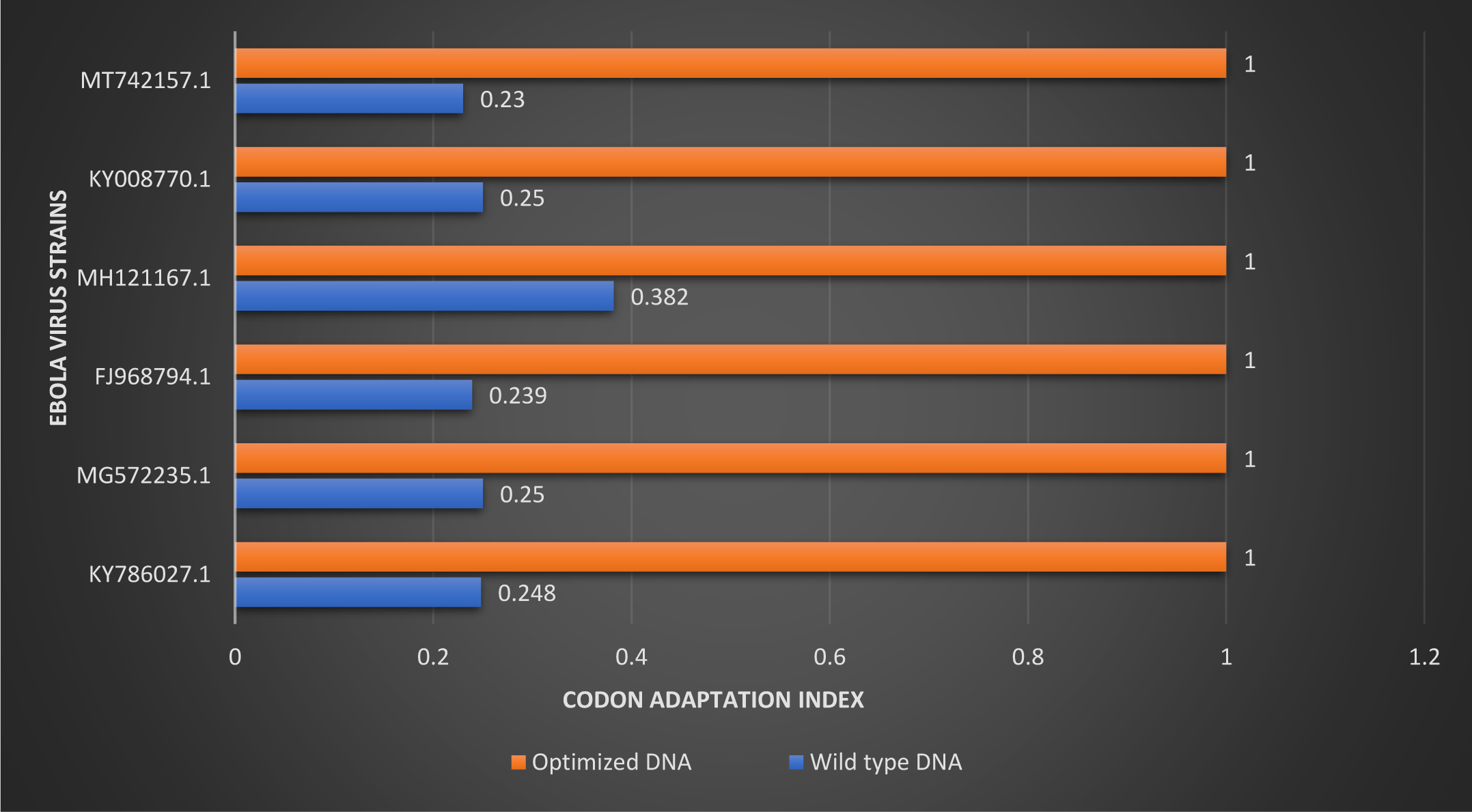
Graph showing comparison between the wild-type and optimized DNA for VP24 gene.

**Fig. 6B:**
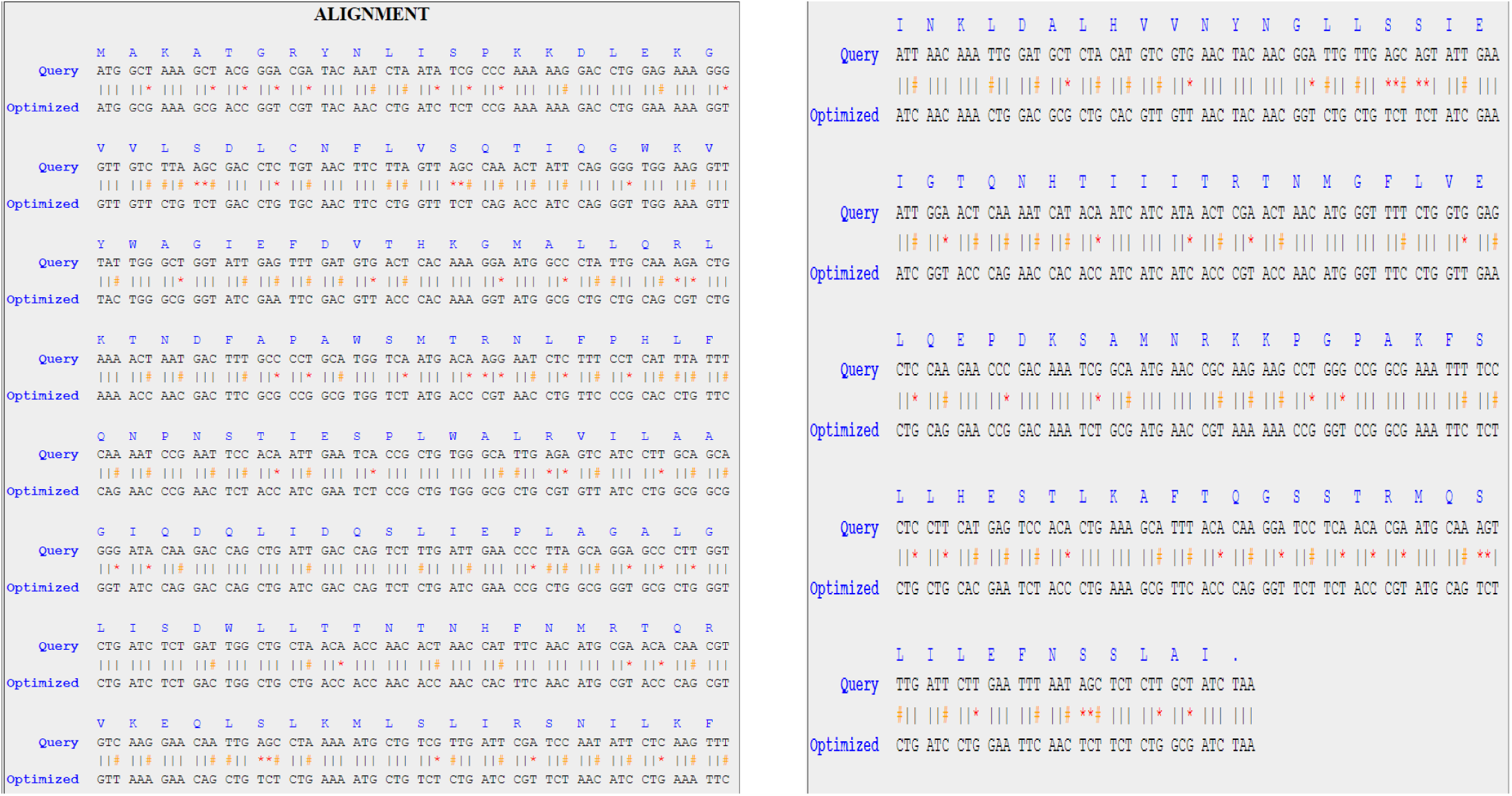
The sequence alignment of wild-type (Query) and codon-optimized DNA for VP24 gene of Ebola virus for accession number MG572235.1. *The specific-coloured symbols stand for (*): Transversion change, (#): Transition change and (|): Unchanged nucleotide

The wild-type L gene exhibited a range of 0.23 to 0.382 for CAI, 39.6 to 40.6 for GC content, and 59.4 to 60.4 for AT content across the six different strains. The average values (±SD) were 0.26 (±0.057) for CAI, 39.98 (±0.386) for GC content, and 60 (±0.430) for AT content (Table 7). Upon codon optimization, the frequencies of GC and AT content in the respective DNA sequences ranged from 32.3 to 53.7 and 46.3 to 67.7, with average values (±SD) of 49.18 (±8.374) and 50.98 (±9.352), respectively. The CAI for the optimized DNA was 1 for all six strains.

**Table 7:**
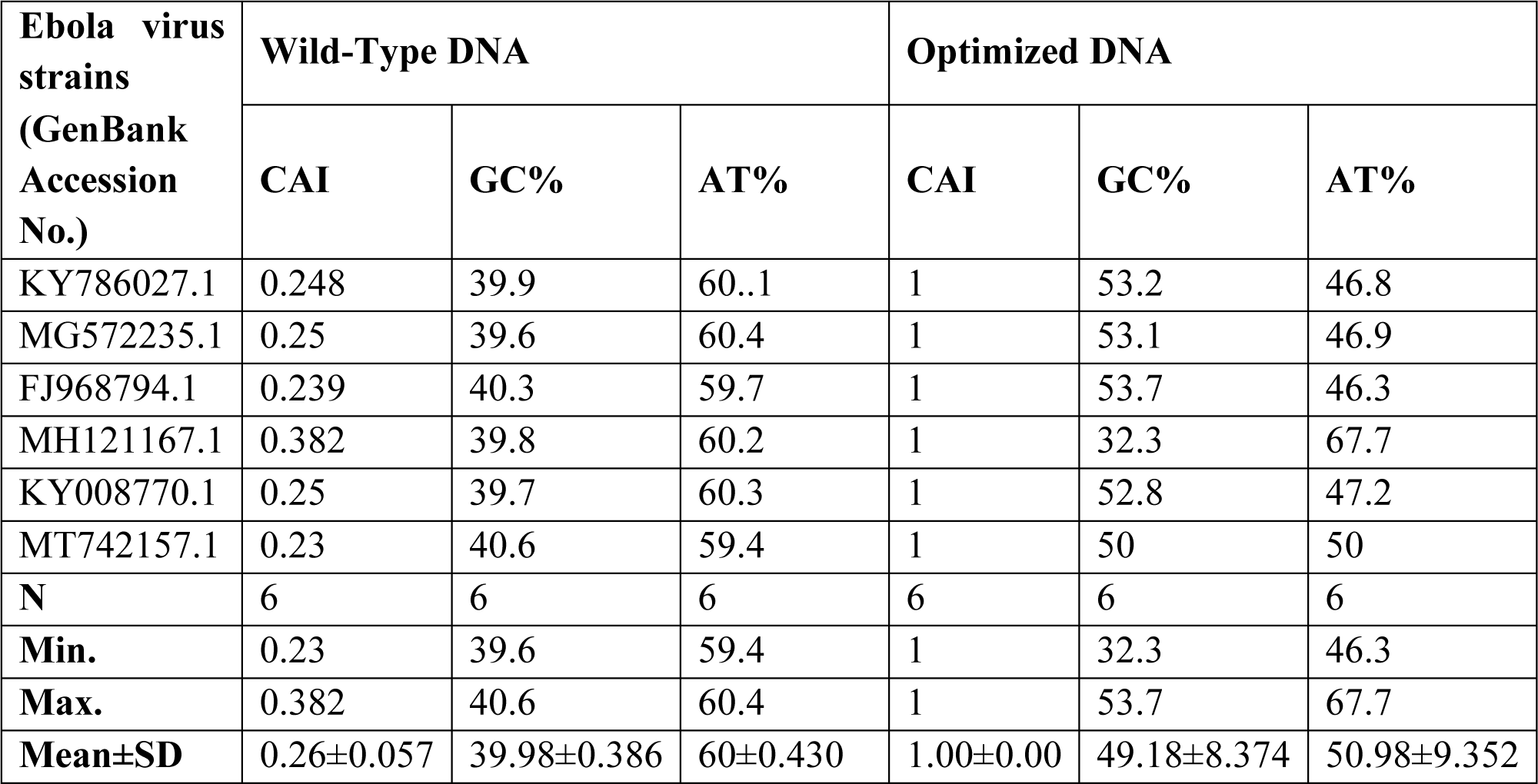
The expression level of polymerase (L) gene of Ebola virus in *E. coli* of wild-type and codon-optimized sequences.

When the mean values of CAI, GC content, and AT content for the polymerase gene in all six strains were compared, the optimized DNA had considerably higher values. The optimized DNA’s mean CAI and GC content were found to be 3.84 (284.61%) and 1.23 (23%), respectively, times higher than the wild-type sequences’ respective mean values. The mean AT content in the optimized DNA, on the other hand, was reduced by 15.03% as compared to the wild-type sequences (Table 7). As illustrated in Fig. 7A, a graph reflecting these comparisons was created. As shown in Fig. 7B, the polymerase gene sequences of both the wild-type and codon-optimized sequences were aligned.

**Fig. 7A:**
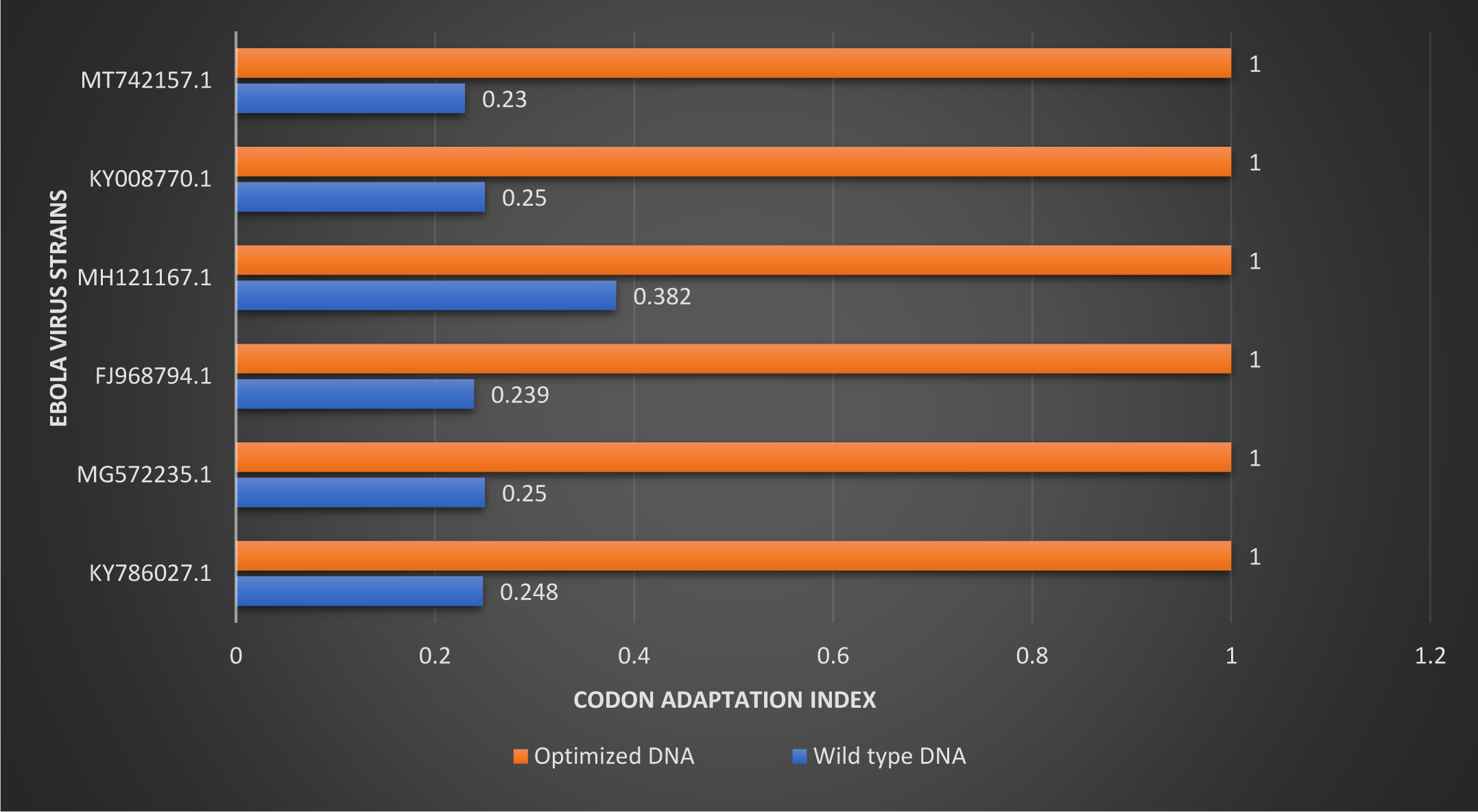
Graph showing comparison between the wild type and optimized DNA for polymerase (L) gene.

**Fig. 7B:**
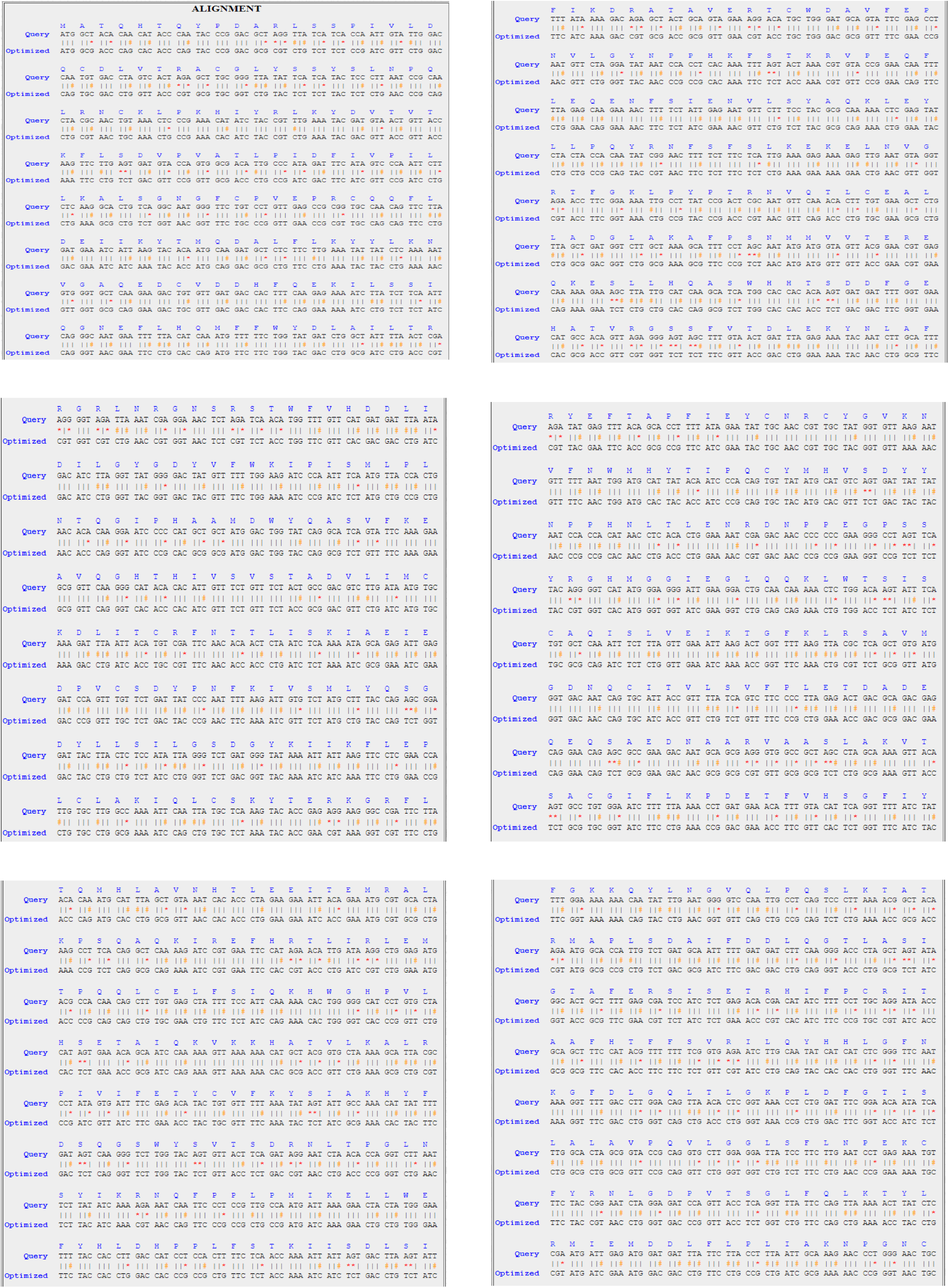
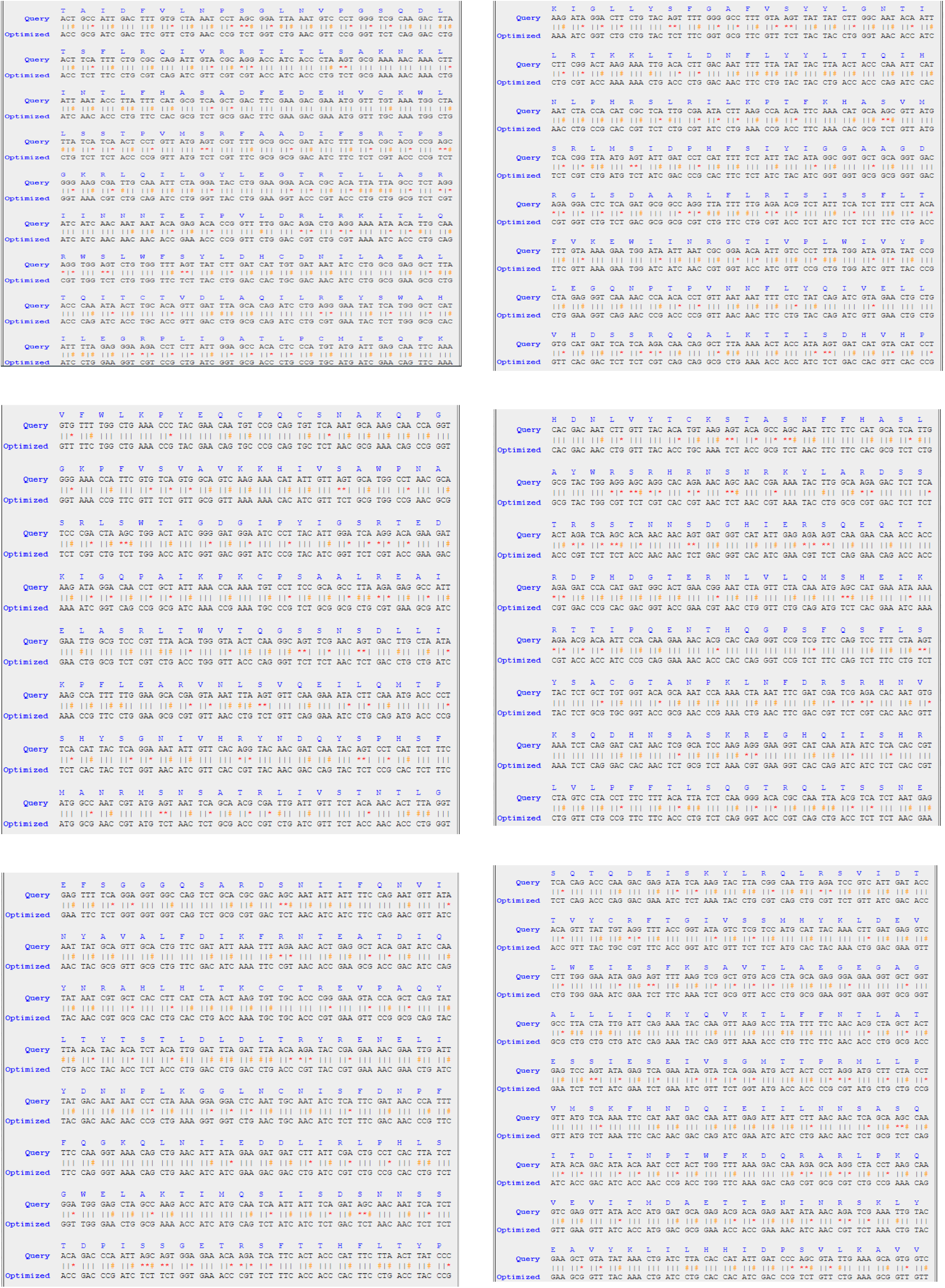
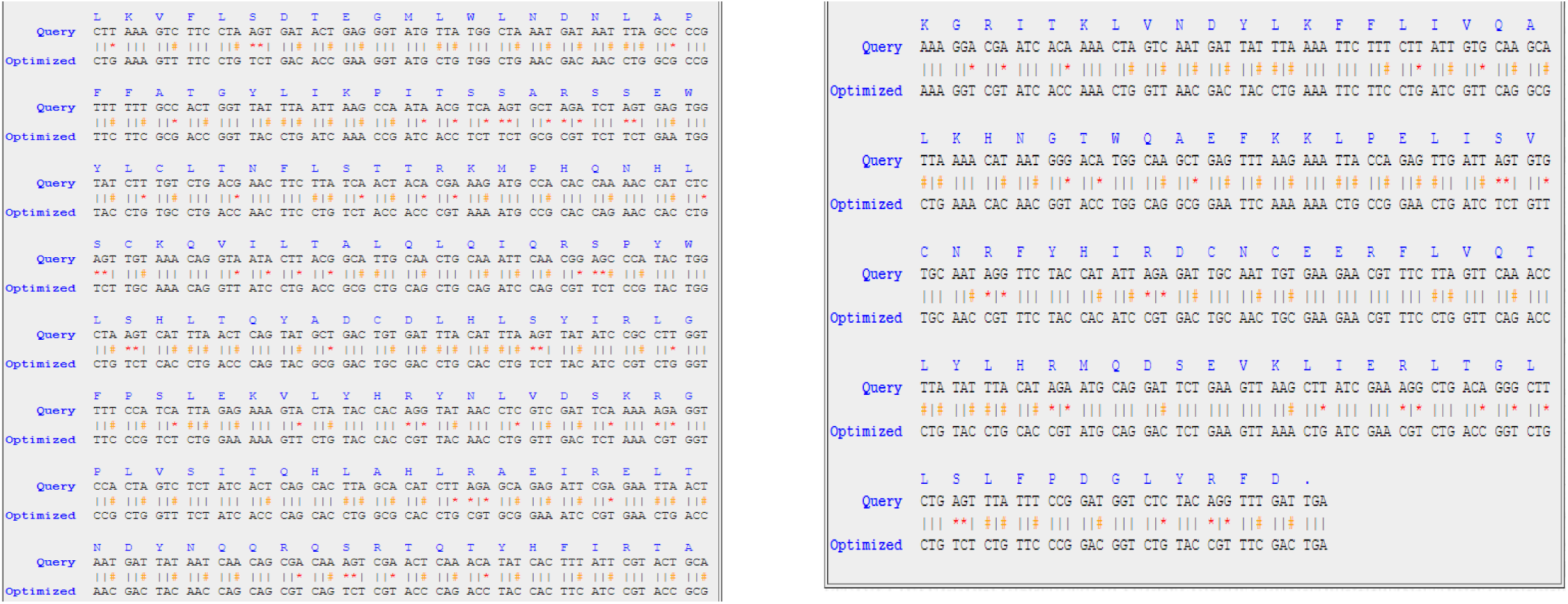
The sequence alignment of wild-type (Query) and codon-optimized DNA for polymerase (L) gene of Ebola virus for accession number MG572235.1. *The specific-coloured symbols stand for (*): Transversion change, (#): Transition change and (|): Unchanged nucleotide.

## 4. Discussions

Codon optimization is a widely used technique aimed at maximizing the expression of protein by addressing limitations associated with the use of codon. This approach involves modifying the DNA sequence to optimize codon selection, which can enhance gene functionality, improve protein expression levels, lower production costs, and facilitate drug development. Codon optimization finds application in various contexts, from animal testing to removing stop codons, to improving gene functionality and protein expression levels. The manufacture of therapeutic antibodies, cytokines, and fusion proteins has considerably increased patient outcomes in the field of biotherapeutics, which plays a crucial role in the treatment of numerous human illnesses, notably in cancer and hematology. Codon optimization is a common expression technique used to improve protein levels. This is feasible due to the genetic code’s degeneracy, which permits most amino acids to be encoded by numerous synonymous codons. The choice of codons can have a significant impact on the protein expression levels. As a result, codon optimization is widely employed in bioproduction and *in vivo* nucleic acid medicinal applications. Studies have shown that codon optimization can significantly increase protein expression, with some reports indicating over a 1000-fold improvement (Gustafsson et al., 2012). However, most studies demonstrate more modest increases in protein expression levels. Surprisingly, synonymous codon mutations have also been used to fine-tune expression through de-optimization. In the case of bispecific antibodies, for example, de-optimization of one of the light chain genes resulted in enhanced antibody production (Magistrelli et al., 2017). DNA-based immunity has shown promise in human trials for HIV infection. In one study, adapting the codon use of the HIV gag protein given by a DNA vaccination resulted in a 10-fold increase in gene expression compared to the wild-type sequence (Menzella 2011). Gene optimization has also been shown to be beneficial in a variety of therapy applications where a protein is synthesized *in vivo* after gene delivery. This method is currently widely employed in a variety of applications (Puigbo et al., 2007).

Codon optimization has been demonstrated to enhance the production of beta-glucanase by *Streptomyces* spp. genes when expressed in heterologous hosts. This optimization process has shown profitable results in terms of increased protein yield and improved functional activity. Optimized and synthesized genes perform better than their wild-type counterparts due to their influence on mRNA stability and proper protein folding, leading to increased enzyme activity. Gene optimization influences several molecular processes, including transcription and translation, resulting in increased product yield and activity (Edison et al., 2020). Codon optimization also has the advantage of preventing stable hairpin formation at the 5’ mRNA end, facilitating optimal mRNA loading and protein translation. Research including the development of 154 GFP mutants in *E. coli* found that constructed hairpins at the 5’ mRNA end drastically lowered GFP expression by up to 250-fold when compared to an adequately codon-optimized construct (Sander et al., 2014). In addition to enhancing protein expression levels, codon usage analysis has found applications in metagenomic studies. For example, codon usage frequency analysis has been utilized to determine the host range of RNA virus genomes from high temperature acidic metagenomes, whether they originate from bacteria, archaea, or eukaryal (Stedman et al., 2013).

In the present study, we calculated mean values of GC content, CAI, and AT content for all EBOV strains. We compared these values with the corresponding values obtained for the optimized DNA sequences. The results concluded that the sequences of optimized DNA had noticeably different and greater values compared to their respective wild-type strains for all genes. The CAI values of the optimized DNA sequences were found to be considerably higher when analyzing the NP gene of all six strains. In the optimized DNA, the CAI of the NP gene was increased by 3.14-fold (213.5%). Similarly, the CAI values in the optimized DNA were enhanced by 3.44-fold (244.8% increase) for the VP35 gene, 84-fold (284.6% increase) for the VP40 gene, and 57-fold (257.14% increase) for the GP gene. The VP30 gene showed an increase of 34-fold (334.8% increase), while the VP24 gene showed an increase of 84-fold (284.61% increase). Additionally, the L gene exhibited an increase of 3.84-fold (284.61% increase).

These findings indicate that the optimization of DNA sequences led to significantly higher CAI values for various EBOV genes, indicating improved codon usage and potentially enhanced gene expression. The study also discovered an increase in the proportion of GC content in optimized DNA sequences as compared to wild-type sequences. The GC content of the NP gene in the optimized DNA was increased by 1.2-fold (19.17% increase) on average. Similarly, the GC content in the optimized DNA of VP35 and VP40 increased by 1.22-fold (22.5% increase) and 1.2-fold (21.2% increase), respectively. The VP24 gene showed an increase of 1.23-fold (23.3% increase), and the L gene demonstrate an increase of 1.23-fold (23% increase). Still, the AT content in all genes was considerably lower as compared to wild-type sequences. Similarly in silico codon optimization approach has been utilized in various microorganisms such as *Mycobacterium tuberculosis* (Mani et al., 2010), *Aeromonas hydrophila* (Singh et al., 2010), influenza A virus (Mani et al., 2011), SARS-COV-2 (Al Zamane et al., 2021), Nipah virus (Gupta et al., 2022), etc.

EBOV survivors’ humoral responses largely target the surface GP, and the existence of anti-GP neutralising antibodies has been linked to protection against EBOV infection. The EBOV surface glycoproteins GP1, 2 are important for host cell adhesion and fusion and are the principal target of neutralising antibodies. High levels of GP1, 2 expressions can disturb normal cell physiology, and EBOV regulates GP gene expression via an RNA-editing process. Codon optimization can be used to increase GP1, 2 expressions, which reduces the synthesis and release of EBOV virus-like particles (VLPs) as well as the infectivity of GP1, 2-viruses of the pseudo-type. This impact is mediated by two different processes. For starters, increased GP1, 2 expression reduces the synthesis of other proteins essential for viral assembly. Second, viruses with high amounts of GP1, 2 are fundamentally less infectious, which might be owing to poor receptor binding or endosomal processing (Mohan et al., 2015). These findings suggest that the expression levels of GP1, 2 have a significant impact on factors contributing to virus fitness and that RNA editing may play a crucial role in regulating GP1, 2 expressions to optimize virus production and infectivity.

Several vaccine candidates are currently being developed to prevent EVD by utilizing the Ebola virus GP and NP to elicit a protective immune response in preclinical animal models. Two vaccine strategies that have shown promise in preventing EVD include: Recombinant vesicular stomatitis virus (rVSV)-based vaccine, which uses a modified vesicular stomatitis virus (VSV) vector to express the Zaire ebolavirus (ZEBOV) surface GP. The vaccination is called rVSVG-ZEBOV-GP. It has been used in preclinical research and has been granted WHO prequalification. This vaccine has demonstrated effectiveness in preventing EBOV infection. Adenovirus and modified vaccinia ankara (MVA) vector-based vaccine: Adenovirus type 26-vectored vaccine encoding the ZEBOV glycoprotein (Ad26.ZEBOV) is used in this vaccination method, followed by a booster dose of a modified vaccinia Ankara virus strain (MVA-BN-Filo). This sequential immunisation strategy has also gained WHO prequalification status and has demonstrated promising effects in EVD prevention (Henao et al., 2017; Gsell et al., 2016; Anywaine et al., 2019; Mutua et al., 2019; Barry et al., 2021; Afolabi et al., 2022; Ishola et al., 2022;). These vaccine candidates have been used in response to recent Ebola outbreaks and are considered important tools in controlling and preventing the spread of the EBOV. In clinical studies done by the Partnership for Research on Ebola Vaccinations (PREVAC) collaboration, the rVSVG-ZEBOV-GP vaccine and the Ad26.ZEBOV-MVA-BN-Filo combo showed encouraging outcomes. The rVSVG-ZEBOV-GP vaccine is a one-dose vaccine that is suggested for reactive ring vaccination in people who are at high risk of Ebola virus infection during epidemics. The Ad26.ZEBOV-MVA-BN-Filo combination is a two-dose regimen suggested for those at low to moderate risk of Ebola virus infection (Lévy et al., 2018). GP and NP, known for their immunogenic properties, are suitable vaccine candidates against EBOV.

Codon optimization can be used to boost gene expression in cells, providing high titers for a large-scale vaccine manufacture. By optimizing codons, protein expression levels can be improved, leading to more effective vaccines. The combination of codon optimization and immunology-based studies is crucial for the development of effective vaccines against EBOV. This approach enables the production of vaccines with enhanced immunogenicity and potency, which can be upscaled for industrial production. Developing and manufacturing such vaccines might be a significant therapeutic technique for preventing EBOV infection. It is crucial to note, however, that codon optimization may create certain difficulties. It requires a comprehensive understanding of codon usage patterns in the target host or expression system. Additionally, modifying codon usage can potentially affect protein folding, stability, or function, necessitating careful analysis and experimental validation to ensure the desired vaccine characteristics are maintained.

To address these challenges, it is essential to conduct *in vitro* analyses and experimental validation to complement in silico studies. *In vitro* analysis allows for a more comprehensive understanding of the specific protein and its interactions within the biological system, providing insights into the impact of codon optimization on protein structure, function, and post-translational modifications. By conducting thorough *in vitro* studies, researchers can overcome these challenges associated with codon optimization and ensure the safety and efficacy of biotech therapeutics. It is crucial to carefully evaluate the consequences of synonymous codon changes on protein properties to develop effective and reliable biotech products. Moving ahead, it is critical to emphasise awareness of the differences between EBOV species. More field research on reservoir species ecology and shedding techniques is desperately needed. Exploring the pathophysiology of EBOV infections in laboratory animals is crucial for identifying novel targets for intervention strategies. These research activities attempt to produce medications that are both accessible and economical for the treatment of EBOV infections. A thorough and well-defined plan is necessary to successfully convert prospective medications and vaccines from the laboratory to clinical trials and, eventually, to treat Ebola patients. Prevention of disease spread remains the best approach to reduce the number of cases and control epidemics. Large-scale awareness programs should be organized to educate the public and promote disease eradication efforts.

## 5. Conclusions

In conclusion, the development of vaccines based on GP or NP, combined with codon optimization and immunology-based studies, offers great promise in combating EBOV infection. This approach can facilitate the production of effective vaccines at an industrial scale. However, it is essential to address the challenges associated with codon optimization to ensure the safety and efficacy of resulting vaccines.

While there are several applications for codon optimization in the development of recombinant protein therapeutics and nucleic acid treatments, it is vital to examine potential obstacles. Reports suggest that synonymous codon changes introduced during optimization may impact protein conformation, stability, and function, as well as increase immunogenicity. Such alterations could have unintended consequences, including reduced efficacy or association with certain diseases.

The current study focuses on optimizing codons to enhance the expression of different EBOV genes in *E. coli*, overcoming limitations caused by codon bias. The optimized sequences showed increased CAI and GC content, indicating their potential for overexpression in *E. coli*. Based on this study and previous research on immunogenicity, GP or NP are promising candidates for an EBOV vaccine, as they possess immunogenic properties and can stimulate an immune response.

*In vitro* validation of the in silico study’s findings is the next step, assessing the level of overexpression achieved by the optimized sequences and evaluating their safety and efficacy in generating an immune response. Successful validation could lead to further development of these optimized genes on an industrial scale for immunodiagnostic tools and immunotherapeutic. It’s crucial to emphasize that rigorous testing is essential to confirm the findings and ensure the safety and potency of potential vaccines. These efforts will contribute to the development of effective immunodiagnostics and immunotherapeutic strategies to combat EBOV infections, providing hope for controlling this deadly disease.

## Acknowledgments

The authors are thankful to the Principal, of Gargi College for providing the infrastructural support.

## References

1. Afolabi MO, Ishola D, Manno D, et al. Safety and immunogenicity of the two-dose heterologous Ad26.ZEBOV and MVA-BN-Filo Ebola vaccine regimen in children in Sierra Leone: a randomised, double-blind, controlled trial. Lancet Infect Dis 2022;22:110–122.

2. Al Zamane S, Nobel FA, Jebin RA, Amin MB, Somadder PD, Antora NJ, Hossain MI, Islam MJ, Ahmed K, Moni MA. Development of an in silico multi-epitope vaccine against SARS-COV-2 by précised immune-informatics approaches. Inform Med Unlocked. 2021;27:100781. doi: 10.1016/j.imu.2021.100781.

3. Aleksandrowicz P., Marzi A., Biedenkopf N., et al.: Ebola virus enters host cells by macropinocytosis and clathrin-mediated endocytosis. J Infect Dis. 2011; 204 Suppl 3: S957–967.

4. Anywaine Z, Whitworth H, Kaleebu P, et al. Safety and immunogenicity of a 2-dose heterologous vaccination regimen with Ad26.ZEBOV and MVA-BN-Filo Ebola vaccines: 12-month data from a phase 1 randomized clinical trial in Uganda and Tanzania. J Infect Dis 2019;220:46-56.

5. Ascenzi P., Bocedi A., Heptonstall J. et al.: Ebolavirus and Marburgvirus: insight the Filoviridae family. Mol Aspects Med. 2008; 29: 151–185.

6. Barry H, Mutua G, Kibuuka H, et al. Safety and immunogenicity of 2-dose heterologous Ad26.ZEBOV, MVA-BN-Filo Ebola vaccination in healthy and HIV-infected adults: a randomised, placebo-controlled phase II clinical trial in Africa. PLoS Med 2021;18(10):e1003813–e1003813.

7. Biedenkopf N., Hartlieb B., Hoenen T. et al.: Phosphorylation of Ebola virus VP30 influences the composition of the viral nucleocapsid complex: impact on viral transcription and replication. J Biol Chem. 2013; 288: 11165–11174.

8. Carroll MW, et al. Deep sequencing of RNA from blood and oral swab samples reveals the presence of nucleic acid from a number of pathogens in patients with acute Ebola virus disease and is consistent with bacterial translocation across the gut. mSphere. 2017;2:e00325–17. doi: 10.1128/mSphereDirect.00325-17.

9. Carroll S.A., Towner J.S., Sealy T.K., et al.: Molecular evolution of viruses of the family Filoviridae based on 97 whole-genome sequences. J Virol. 2013; 87: 2608–2616.

10. Edison L. K, Dan V. M, Reji S. R, Pradeep N. S. A Strategic Production Improvement of Streptomyces Betaglucanase Enzymes with Aid of Codon Optimization and Heterologous Expression. Biosci Biotech Res Asia 2020;17(3):587-599.

11. Feldman H., Sanchez A., Geisbert T.: Filoviridae: Marburg and Ebola viruses. In Fields Virology, sixth edition. Ed. DM Knipe & PM Howley; Lippincott Williams & Wilkins, Wolters Kluwer; Philadelphia 2013, v.2, 923–956.

12. Food and Drug Administration. First FDA-approved vaccine for the prevention of Ebola virus disease, marking a critical milestone in public health preparedness and response. December 19, 2019

13. Geisbert T.W., Jahrling P.B.: Differentiation of filoviruses by electron microscopy. Virus Res. 1995; 39: 129–150.

14. Goeijenbier M., van Kampen J.J., Reusken C.B., Koopmans M.P., van Gorp E.C.: Ebola virus disease: a review on epidemiology, symptoms, treatment and pathogenesis. Neth J Med. 2014; 729: 442–448.

15. Graf M, Schoedl T, Wagner R (2009) Rationals of Gene Design and de novo Gene Construction. Systems Biology and Synthetic Biology pp 411–438

16. Gsell P-S, Camacho A, Kucharski AJ, et al. Ring vaccination with rVSV-ZEBOV under expanded access in response to an outbreak of Ebola virus disease in Guinea, 2016: an operational and vaccine safety report. Lancet Infect Dis 2017;17:1276-1284.

17. Gupta A, Gangotia D, Vasdev K and Mani I (2022). In silico DNA codon optimization of the variant antigen-encoding genes of diverse strains of Nipah virus. Indian J. Biotech. Pharm. Res. 10(1):1–16.

18. Gustafsson C, Minshull J, Govindarajan S, Ness J, Villalobos A, Welch M. Engineering genes for predictable protein expression. Protein Expr Purif. 2012;83(1):37–46.

19. Gustafsson, C, Govindarajan S, Minshull J (2004) Codon bias and heterologous protein expression. Trends Biotechnol 22:346–353

20. Heinzelman P, Snow CD, Wu I, Nguyen C, Villalobos A, Govindarajan S et al (2009) A family of thermostable fungal cellulases created by structure-guided recombination. P Natl Acad Sci USA 106: 5610–5615. doi: 10.1073/pnas.0901417106

21. Henao-Restrepo AM, Camacho A, Longini IM, et al. Efficacy and effectiveness of an rVSV-vectored vaccine in preventing Ebola virus disease: final results from the Guinea ring vaccination, open-label, cluster-randomised trial (Ebola Ça Suffit!). Lancet 2017;389:505–518.

22. Hoenen T., Groseth A., Kolesnikova L. et al.: Infection of naive target cells with virus-like particles: implications for the function of Ebola virus VP24. J Virol. 2006 Jul; 80 (14): 7260– 7264.

23. ICTV Virus Taxonomy 2013: http://www.ictvonline.org/virusTaxonomy.asp.

24. Ikemura T (1981) Correlation between the abundance of Escherichia coli transfer RNAs and the occurrence of the respective codons in its protein genes. J Mol Biol 146:1–21

25. Ishola D, Manno D, Afolabi MO, et al. Safety and long-term immunogenicity of the two-dose heterologous Ad26.ZEBOV and MVA-BN-Filo Ebola vaccine regimen in adults in Sierra Leone: a combined open-label, non-randomised stage 1, and a randomised, double-blind, controlled stage 2 trial. Lancet Infect Dis 2022;22:97-109.

26. Jain S, Martynova E, Rizvanov A, Khaiboullina S, Baranwal M. Structural and Functional Aspects of Ebola Virus Proteins. Pathogens. 2021 Oct 15;10(10):1330. doi: 10.3390/pathogens10101330. PMID: 34684279; PMCID: PMC8538763.

27. Kanapathipillai R, Henao Restrepo AM, Fast P, Wood D, Dye C, Kieny MP, Moorthy V. Ebola vaccine--an urgent international priority. N Engl J Med. 2014 Dec 11;371(24):2249–51. doi: 10.1056/NEJMp1412166

28. Lévy Y, Lane C, Piot P, et al. Prevention of Ebola virus disease through vaccination: where we are in 2018. Lancet 2018; 392:787–790.

29. Magistrelli G, Poitevin Y, Schlosser F, Pontini G, Malinge P, Josserand S, et al. Optimizing assembly and production of native bispecific antibodies by codon de-optimization. mAbs. 2017;9(2):231–9.

30. Mani I, Chaudhary DK, Somvanshi P, and Singh V (2010). Codon optimization of the potential antigens encoding genes from *Mycobacterium tuberculosis*. Int J Appl Biol Pharm I(2): 292–301.

31. Mani I, Singh V, Chaudhary DK, Somvanshi P, and Negi MPS (2011). Codon optimization of the major antigen encoding genes of diverse strains of influenza A virus. Interdiscip Sci. 3: 1–7. (IF-3.492).

32. Mateo M., Carbonnelle C., Martinez M.J. et al.: Knockdown of Ebola virus VP24 impairs viral nucleocapsid assembly and prevents virus replication. J Infect Dis. 2011; 204 Suppl 3: S892–896.

33. Menzella HG (2011) Comparison of two codon optimization strategies to enhance recombinant protein production in Escherichia coli. Microb Cell Fact10:15. doi: 10.1186/1475-2859-10-15

34. Merck. ERVEBO [Ebola Zaire vaccine (rVSVΔG-ZEBOV-GP) live] awarded prequalification status by the World Health Organization (WHO). November 13, 2019

35. Mohan GS, Ye L, Li W, Monteiro A, Lin X, Sapkota B, Pollack BP, Compans RW, Yang C. Less is more: Ebola virus surface glycoprotein expression levels regulate virus production and infectivity. J Virol. 2015 Jan 15;89(2):1205–17. doi: 10.1128/JVI.01810-14. Epub 2014 Nov 12. PMID: 25392212; PMCID: PMC4300637.

36. Mutua G, Anzala O, Luhn K, et al. Safety and immunogenicity of a 2-dose heterologous vaccine regimen with Ad26.ZEBOV and MVA-BN-Filo Ebola vaccines: 12-month data from a phase 1 randomized clinical trial in Nairobi, Kenya. J Infect Dis 2019; 220:57-67.

37. Nouvellet P, et al. The role of rapid diagnostics in managing Ebola epidemics. Nature. 2015;528:S109–S116. doi: 10.1038/nature16041.

38. Puigbo P, Guzmen E, Romeu A, Garcia-Vallve S (2007) OPTIMIZER: A web server for optimizing the codon usage of DNA sequences. Nucleic Acids Res 35:W126–W131

39. Sander IM, Chaney JL, Clark PL (2014) Expanding Anfinsen’s principle: contributions of synonymous codon selection to rational protein design. J Am Chem Soc 136(3):858–61.

40. Singh V, Chaudhary DK, Mani I, Somvanshi P, Rathore G, and Sood N (2010). Molecular Identification and codon optimization analysis of major virulence encoding genes of *Aeromonas hydrophila*. African Journal of Microbiology Research. 4(10): 952–957.

41. Stahelin R.V.: Membrane binding and bending in Ebola VP40 assembly and egress. Front Microbiol. 2014; 5: 300. doi: 10.3389/fmicb.2014.00300. eCollection 2014.

42. Stedman KM, Kosmicki NR, Diemer GS (2013) Codon usage frequency of RNA virus genomes from high-temperature acidic-environment metagenomes. J Virol 87(3):1919.

43. Tomori O, Kolawole MO. Ebola virus disease: current vaccine solutions. Curr Opin Immunol. 2021 Aug;71:27–33. doi: 10.1016/j.coi.2021.03.008.

44. Towner J.S., Amman B.R., Sealy T.K., et al.: Isolation of genetically diverse Marburg viruses from Egyptian fruit bats. PLoS Pathog. (2009), 5:e1000536.

45. Vernet M-A, et al. Clinical, virological, and biological parameters associated with outcomes of Ebola virus infection in Macenta, Guinea. JCI Insight. 2017;2:e88864. doi: 10.1172/jci.insight.88864.

46. Wannier SR, et al. Estimating the impact of violent events on transmission in Ebola virus disease outbreak, Democratic Republic of the Congo, 2018– 2019. Epidemics. 2019;28:100353. doi: 10.1016/j.epidem.2019.100353.

47. World Health Organization. Urgently Needed: Rapid, Sensitive, Safe and Simple Ebola Diagnostic Tests. http://www.who.int/mediacentre/news/ebola/18-november-2014-diagnostics/en (2014).

